# Do egg hormones have fitness consequences in wild birds? A systematic review and meta-analysis

**DOI:** 10.1101/2024.10.29.620852

**Authors:** Lucia Mentesana, Michaela Hau, Pietro B. D’Amelio, Nicolas M. Adreani, Alfredo Sánchez-Tójar

**Author notes:** Author for correspondence: Dr. Lucia Mentesana, Address: Igua 4225, Montevideo (11400), Uruguay. Dr. Michaela Hau, Dr. Pietro B. D’Amelio, Dr. Nicolas M. Adreani, Dr. Alfredo Sánchez-Tójar.

## Abstract

Egg-laying species are key models for understanding the adaptive significance of maternal effects, with egg hormones proposed as an important underlying mechanism. However, even thirty years after their discovery, the evolutionary consequences of hormone-mediated maternal effects remain unclear. Using evidence synthesis, we tested the extent to which increased prenatal maternal hormone deposition in eggs relates to fitness in wild birds (19 species, 438 effect sizes, 57 studies). Egg androgens, glucocorticoids, and thyroid hormones showed an overall near-zero mean effect for both maternal and offspring fitness proxies. However, heterogeneity was high, suggesting that egg hormone effects on fitness are context-dependent. Hormone type and age did not explain much of the observed variance, nor did methodological factors such as type of study or experimental design. Heterogeneity decomposition showed that differences in effect sizes were mostly driven by within-study variability and phylogenetic relationships. Our study provides the most comprehensive investigation to date of the relationship between egg hormones and fitness in vertebrates. By synthesizing current knowledge, we aim to overcome theoretical shortcomings in the field of maternal effects via egg hormone deposition and inspire new research into its many intriguing aspects.

## 1. Introduction

Maternal (genetic and nongenetic) effects are widespread across nature, occurring in plants, invertebrates, and vertebrates. They occur when mothers shape their offspring’s phenotype by influencing the environment they provide (Mousseau & Fox 1998; Reinhold 2002; Uller 2008, 2012; Wolf & Wade 2016). The general contribution of maternal effects to offspring phenotypic variation within populations was found to be half that (∼ 11%) of additive genetic effects (∼ 21%), influencing offspring morphology, physiology, and phenology, among other traits (meta-analysis by Moore *et al*. 2019; but see also Sánchez-Tójar *et al*. 2020; Yin *et al*. 2019). Mothers can exert these effects both before and after birth, but prenatal maternal effects are generally expected to have larger and more irreversible impacts on offspring than postnatal effects, with consequences for offspring survival and reproductive success (Groothuis *et al*. 2005a; Marshall & Uller 2007; Mousseau & Fox 1998; Uller 2008; Wolf *et al*. 2011).

Egg-laying species are key models for understanding the adaptive significance of maternal effects and their underlying mechanisms. Unlike in most mammals, embryos develop outside the mother’s body, limiting the time for direct maternal influence on the offspring. In egg-laying species, mothers can influence the offspring’s prenatal environment by depositing compounds into the egg (e.g., Lovern & Wade 2001; McCormick 1999; Mentesana *et al*. 2019; Valcu *et al*. 2019; Weiss *et al*. 2011). In 1993, Schwabl first reported maternal hormones in bird eggs. These findings then were extended to other egg-laying vertebrates like fishes, reptiles, and amphibians (e.g., Groothuis *et al*. 2005a). Since then, many studies have suggested that hormones, especially steroid hormones present in the yolk, shape offspring growth, behavior (e.g., begging, aggression, boldness, activity, food competitiveness), and physiological systems (e.g., immune system, hypothalamic-pituitary-adrenal axis, oxidative condition; von Engelhardt & Groothuis 2011; Gil 2008; Groothuis *et al*. 2005a; Lovern & Wade 2001; Mentesana *et al*. 2021a). Yet, whether such phenotypic changes translate into fitness benefits for the mother, the father, or the offspring remains under debate (reviewed by Bebbington & Groothuis 2021), especially for wild populations (e.g., Tschirren *et al*. 2014).

Egg hormone-mediated maternal effects are often considered an adaptive mechanism, where mothers are assumed to adjust hormone concentrations in response to environmental cues to shape offspring phenotypes and optimize maternal and/or offspring fitness in variable but predictable environments (reviewed by Groothuis *et al*. 2005a). However, maternal egg hormone concentrations may also lack adaptive value and instead reflect unavoidable physiological processes, such as changes in maternal hormone concentrations that passively diffuse into the egg during formation, leading to varying hormone allocation (reviewed by Gil 2008; Groothuis & Schwabl 2008). Thirty years after the discovery of maternal hormones in eggs, the evolutionary consequences of hormone-mediated maternal effects remain unclear.

There are several reasons why the evolutionary consequences of maternal egg hormones remain unresolved. First, the influence of maternal hormones on fitness is expected to vary greatly, depending not only on the group of hormones studied but also on the fitness proxy used, which can result in contrasting fitness effects. For example, egg androgens (i.e., androstenedione, testosterone, and 5-α dihydrotestosterone) and thyroid hormones (i.e., TH3 and TH4) can lead to increased behavioral competitiveness, boost early growth, and ultimately survival (von Engelhardt & Groothuis 2011; Groothuis *et al*. 2005a; Hsu *et al*. 2017; Ruuskanen *et al*. 2018). However, high concentrations of egg androgens can also impair the offspring’s immune system (reviewed by Gil 2008) and oxidative condition (e.g., Treidel *et al*. 2013), potentially decreasing survival. Regarding egg glucocorticoids, they have been reported to alter behaviors such as antipredator response, impair physiological systems beneficial for offspring (e.g., immune system), and decrease offspring body condition, growth rate, and ultimately survival (Haussmann *et al*. 2012; Polich *et al*. 2018; Stier *et al*. 2009). Second, egg hormones might have sex-specific effects on offspring fitness. A meta-analysis of bird studies investigating how hormonal manipulations of eggs affect offspring traits showed that experimentally increasing androgen and corticosterone concentrations via egg injection tended to lead to larger effect sizes in male compared to female offspring (Podmokła *et al*. 2018). However, the generality of this finding is debated, as the study could only examine the relationship between experimentally injected eggs and offspring fitness proxies in seven populations of wild birds - six of which provided data only on androgens, and one on corticosterone. Third, the influence of maternal hormones on fitness might be dependent on ontogenetic state. Maternal effects are expected and have been observed to weaken throughout offspring ontogeny; however, a meta-analysis examining the contribution of additive genetic phenotypic variation determined by maternal effects, found only partial support for this pattern (Moore *et al*. 2019). Similarly, another meta-analysis that considered parental effects more broadly reported inconclusive evidence for such a weakening effect (Sánchez-Tójar *et al*. 2020). Fourth, theory predicts that the evolution of maternal effects generally, and of hormone allocation specifically, should be shaped by conflicting fitness optima between offspring, mothers, and fathers (Bebbington & Groothuis 2021; Marshall & Uller 2007). For example, high concentrations of egg androgens might increase maternal reproductive success and offspring survival by boosting offspring growth. But at the same time, high concentrations of egg androgens might decrease maternal life expectancy by increasing her workload or even reduce reproductive success if the increased workload cannot be fulfilled through parental care. Thus, if conflicting fitness optima between offspring and mothers occur and are not considered, we expect considerable variation in the net fitness consequences of egg hormone concentrations. Finally, besides biological reasons, methodological aspects might also add uncertainty to our understanding of the evolutionary importance of maternal egg hormones (reviewed by Groothuis *et al*. 2020; Groothuis & Von Engelhardt 2005). A meta-analysis examining the effects of different experimental manipulations of egg androgens and glucocorticoids in wild and captive bird populations provided evidence that factors such as study type (correlational *vs* experimental), manipulation methods (e.g., maternal *vs* egg manipulation), and hormone doses (e.g., within natural range *vs* supra-physiological concentrations) can influence the relationship between egg hormones and fitness proxies (Podmokła *et al*. 2018).

Here, we used an evidence synthesis approach to test the extent to which an increase in prenatal maternal egg hormone concentrations relates to fitness in wild populations of egg-laying vertebrates and to understand the patterns of the effects (i.e., the sources of heterogeneity). We tested six pre-registered predictions, which we divided into biological and methodological categories (pre-registration: Mentesana *et al*. 2021b). By pre-registering our study, we enhanced the distinction between hypothesis testing and hypothesis generation, thereby improving the credibility of our research findings (Ihle *et al*. 2017). Note that we mostly focused on the relationship between egg hormones and offspring fitness rather than parental fitness because the former has received the most empirical attention. Indeed, due to a lack of studies, we could not investigate the relationship between egg hormones and paternal fitness and could only conduct an exploratory analysis for egg hormones and maternal fitness.

### 1. Biological predictions

BP.1. Before testing our pre-registered predictions for each group of hormones, we first calculated the overall relationship between egg hormones and offspring fitness proxies, regardless of hormone group, and estimated heterogeneity across effect sizes. Next, for each hormone group, we based our predictions on the prevailing hypothesis in the literature. We predicted a positive association between egg androgens (BP.1.1) and thyroid hormones (BP.1.3) with offspring fitness proxies, and a negative one with egg glucocorticoids (BP.1.2). We did not have directional predictions for progesterone and estradiol (BP.1.4 & BP.1.5) due to insufficient data (reviewed by Gil 2008; Groothuis *et al*. 2019). For the remaining predictions, we assumed the same hormone-specific directional predictions as detailed in BP.1.

BP.2. Given some early evidence that androgens and glucocorticoids may have a stronger impact on males than females (Podmokła *et al*. 2018), we predicted larger effect sizes in males. For the remaining hormones (thyroids, progesterone, and estradiol), we did not have directional sex-specific predictions. Unfortunately, the number of studies that have addressed whether egg hormones have sex-specific fitness consequences for the offspring in wild populations was too limited to provide reliable conclusions (Supplementary Information S1).

BP.3. Since there is partial evidence that maternal effects weaken throughout ontogeny (Moore *et al*. 2019), we predicted the relationship between egg hormones and offspring fitness proxies to decline from early to late life stages in both altricial and precocial species, weakening from hatching through the nesting period (excluding hatching day) and into adulthood (i.e., before and after offspring independence; BP.3.1). Also, for altricial species, we predicted that the association would become weaker throughout the nesting phase (BP.3.2).

### 2. Methodological predictions

MP.1. We predicted stronger associations in experimental studies than in correlational ones because manipulations should reduce/eliminate confounding factors and/or because physiological constraints or trade-offs might prevent females from depositing optimal hormone concentrations into the egg. An alternative (non-preregistered) prediction would be that manipulating hormone concentrations might incur fitness costs by shifting concentrations away from adaptive values (e.g., von Engelhardt & Groothuis 2011).

MP.2. Since females can potentially modify the transfer of circulating hormones to their eggs (reviewed by Groothuis & Schwabl 2008), we predicted that the association between egg hormones and fitness proxies would be stronger for direct egg manipulations than for female manipulations.

MP.3. The dose of hormones used may differentially affect offspring fitness. While high doses of hormones within the natural range are expected to result in stronger fitness effects, doses beyond naturally-occurring concentrations may lead to weaker or even opposite effects on fitness (e.g., due to toxicity; Norton & Wira 1977). Following Podmokła *et al*. (2018), we predicted that manipulations within the natural range would lead to stronger fitness effects than supra-physiological concentrations for androgens, but not glucocorticoids, whereas we did not have directional predictions for the remaining hormones (thyroids, progesterone, and estradiol).

## 2. Materials and Methods

### 2.1. Protocol

The study protocol was pre-registered on the Open Science Framework before data extraction began (Mentesana *et al*. 2021b; https://doi.org/10.17605/OSF.IO/KU47W). Our pre-registration specified the purpose of the study, our a priori hypotheses and predictions, the data collection procedures, and the entire analytical plan. Unless stated otherwise, we adhered to these pre-registered plans. Throughout, we follow the Preferred Reporting Items for Systematic reviews and Meta-Analyses in ecology and evolutionary biology (PRISMA-EcoEvo Checklist: Supplementary Figure 1 & Supplementary Information S2; O’Dea *et al*. 2021).

### 2.2. Information sources and search

We conducted a systematic search of the literature from 1993 (i.e., the year when the first paper on maternal hormones was published) until the 22nd of August 2022 in both Web of Science Core Collection and Scopus. The search query was designed to find studies that: (1) tested inter- and/or transgenerational (plasticity) effects; (2) measured or experimentally manipulated egg hormone concentrations; and (3) measured maternal and/or offspring fitness proxies. The full keyword search string with further details is provided in Supplementary Information S3.

### 2.3. Study selection and eligibility criteria

Our searches yielded 2404 records which we deduplicated using the R package revtool v.0.3.0 (Westgate 2018). We screened the titles and abstracts of all records using Rayyan (Ouzzani *et al*. 2016). In total, 178 articles passed the title-and-abstract screening and were subjected to full-text screening (for the decision tree used, see Supplementary Figure 2). We included studies conducted in free-living populations of vertebrates that either experimentally manipulated maternal hormone concentrations during egg laying, affecting hormone deposition into eggs, or measured/manipulated egg hormone concentrations during egg laying, provided they also reported fitness proxies. For both mother and offspring, we considered all fitness components related to their reproduction and/or survival (full list of fitness proxies used in Supplementary Information 4). For the offspring, we also considered morphological information because they are commonly accepted as predictors of post-fledging survival (e.g., meta-analysis by Ronget *et al*. 2018).

After full-text screening, we identified 79 articles meeting our criteria (see PRISMA diagram in Supplementary Figure 3). Three observers (LM, NMA and PBD) evenly performed all the screening and double-screened ∼20% of the records to confirm the reproducibility of the procedure (12% for title-and-abstract screening and 30% for full-text screening). The three observers collectively discussed and resolved conflicting decisions (∼7% conflicted papers).

Although our pre-registered plan aimed to test for the relationship between egg hormones and fitness proxies in all egg-laying vertebrate groups, only 2 out of the 79 potentially suitable studies found were conducted on egg-laying vertebrates other than birds; thus, we decided to focus our study solely on birds.

### 2.4. Data collection and extraction

The data extraction was evenly performed by three observers (LM, NMA and PBD). Two observers (LM and AST) double-checked 36% of the extracted data to confirm the reproducibility of this process, LM double-checked the remaining data in search of potential errors, and both observers discussed and fixed any typos found.

We extracted data from text, tables or figures. For the latter, we used the R package metaDigitise v.1.0.1 (Pick *et al*. 2019). In addition, when necessary, we also obtained raw data directly from supplementary materials and/or from published datasets. Complete data extraction was possible for 22 articles (29% of the dataset). To obtain the missing data from the remaining articles, we contacted 31 authors (54 articles) using a standardized email template (Supplementary Information S5), which allowed us to obtain complete data from an additional 20 articles (37% of the dataset). Eight authors communicated that the data were lost, one author expressed that they were working outside science and did not have time to search for the data, another could not provide the data due to being on sabbatical leave, and five authors did not reply. In total, we obtained complete data from 47 articles and partial data for 10 articles (N = 57); that is, we had to exclude 21 in-principle suitable articles (27%) due to preventable reporting issues (for more discussion, see Hennessy *et al*. 2022). During author correspondence and the review process, one author and one reviewer informed us of two studies that although originally fell outside our systematic search (Duckworth *et al*. 2015: k = 1) or did not fulfill our title-and-abstract screening inclusion/exclusion criteria (Remeš 2011: k = 2) contained data for our meta-analysis and were, therefore, subsequently added to our dataset. For a complete list of studies and extracted variables, see Table 1 and the provided data (https://github.com/ASanchez-Tojar/meta-analysis_egg_hormones_and_fitnes).

**Table 1:**
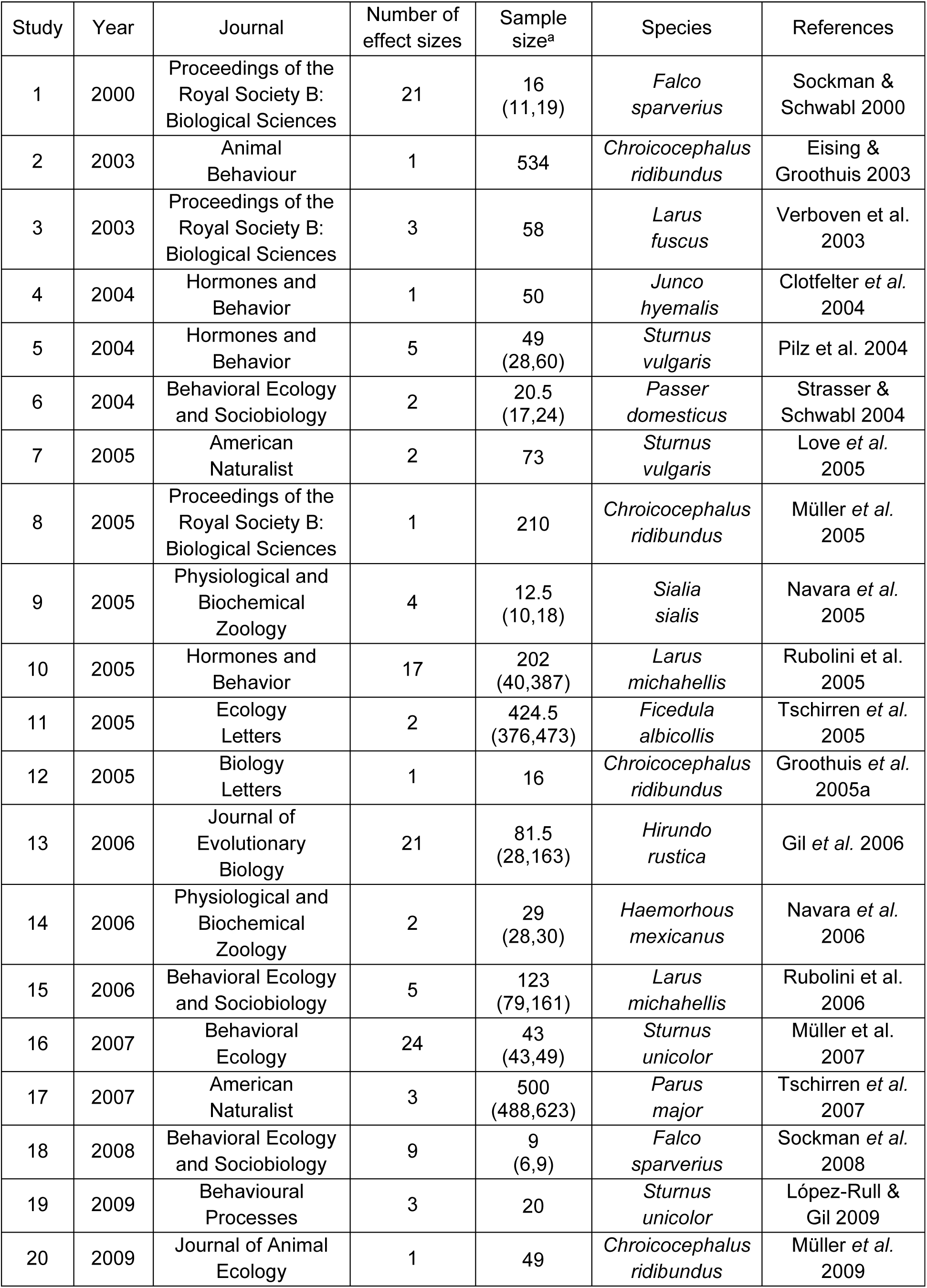
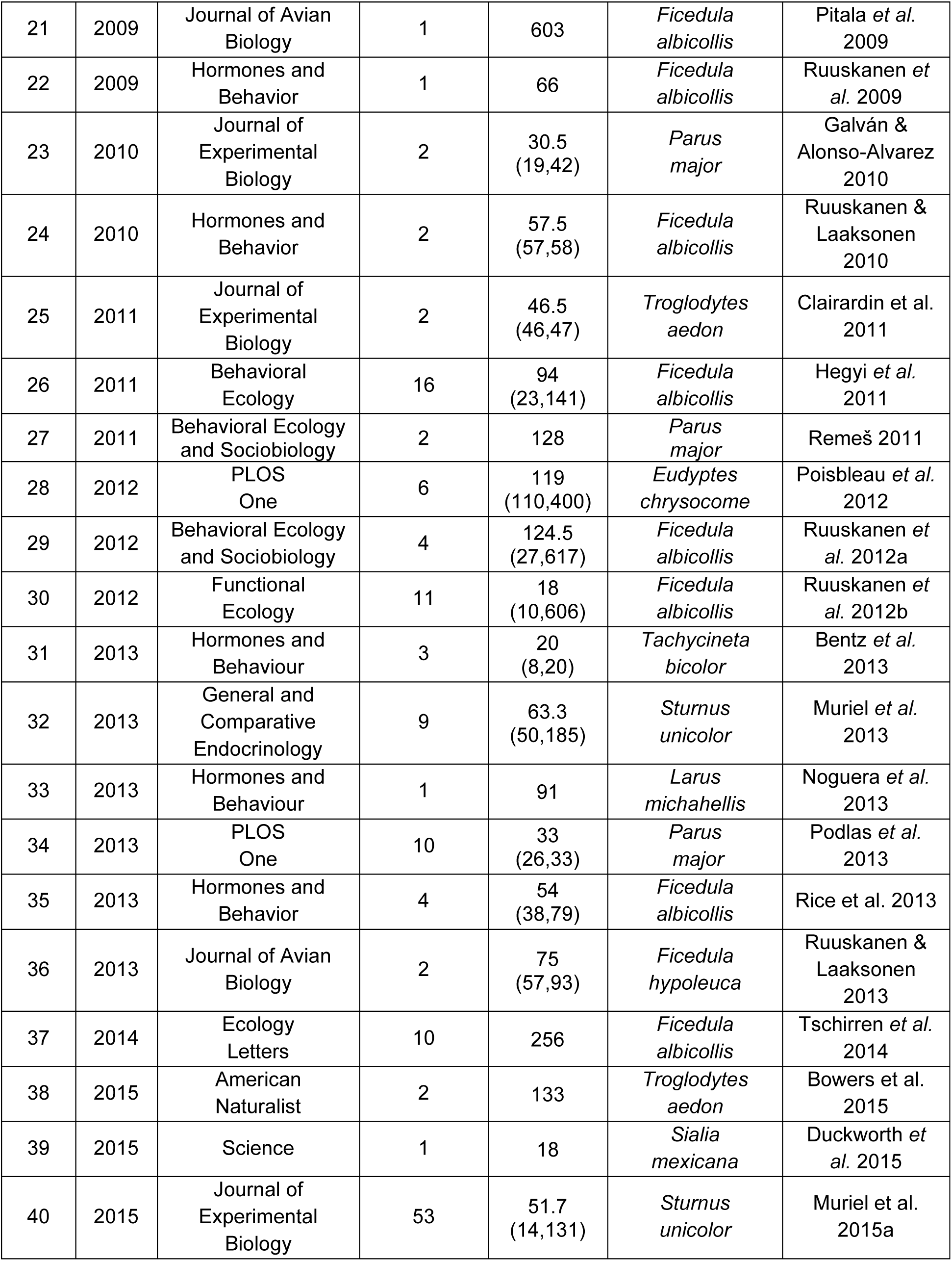
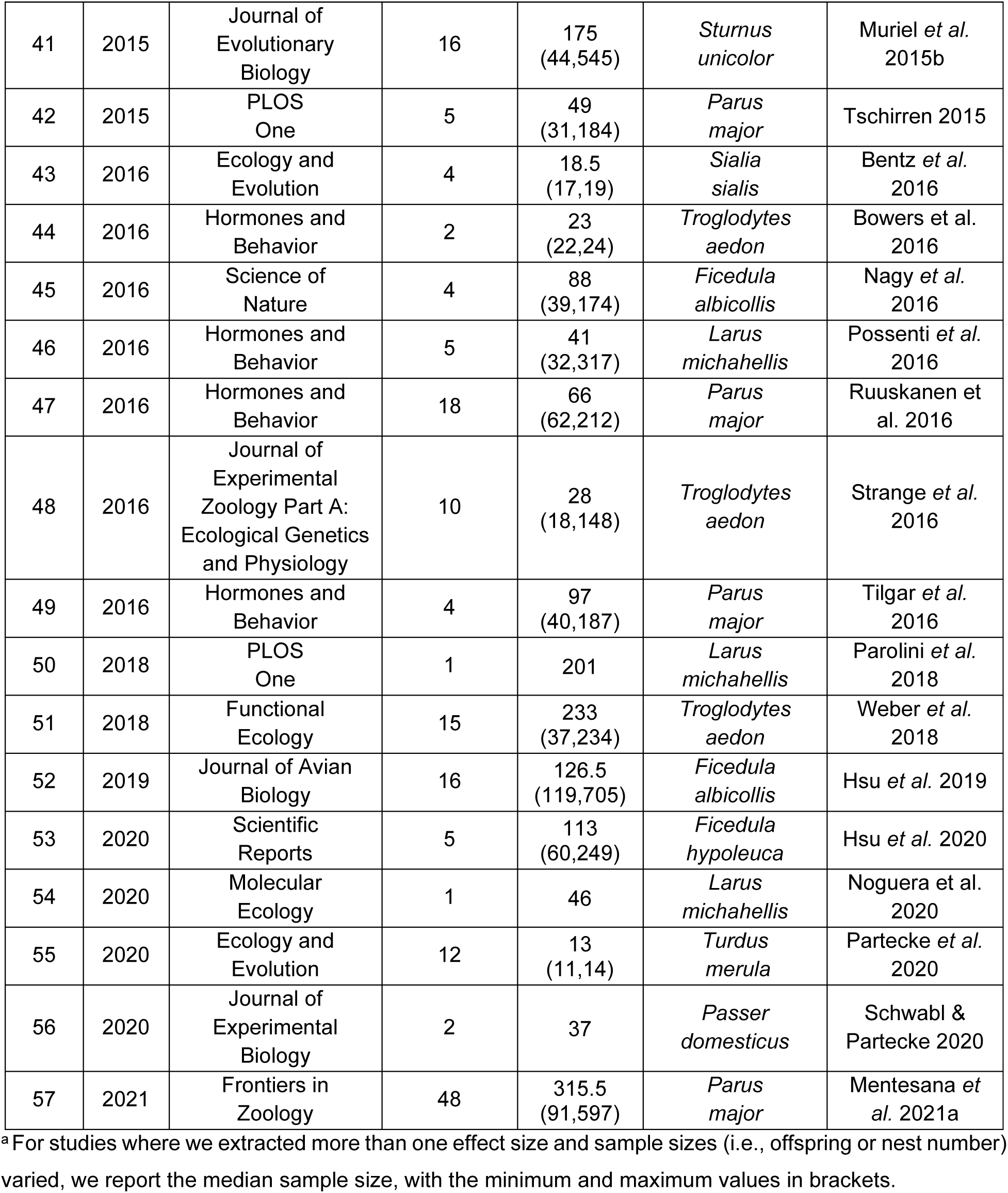
List of studies (N = 57) from which data was extracted and included in our meta-analysis on egg hormone concentrations and maternal and offspring fitness in wild birds.

### 2.5. Extracted variables

From each article we extracted information regarding the study (e.g., title, publication year, author information, journal), the study subjects (e.g., species, developmental mode, mean clutch size), the study area (e.g., location, latitude, longitude and elevation of study area), the methodology (e.g., egg collection, sampling method, subject of the experiment, method and dose use), the fitness proxy studied (e.g., maternal/offspring, trait measured, offspring life history stage), the results (e.g., statistical test used, statistical value, mean values for control and experimental groups), and information on how data were provided (e.g., data location, data source). For the complete list of continuous and categorical moderators see Supplementary Information S6.

Note that for some fitness proxies, it is difficult to fully disentangle maternal and offspring fitness (e.g., hatching or fledging success; discussed by Wolf & Wade 2001). Following the recommendations of Wolf & Wade (2001), we took a conservative approach: fitness proxies measured in offspring that are largely or entirely controlled by the mother (e.g., hatching success, which is strongly influenced by maternal factors such as egg size; meta-analysis by Krist 2011) were assigned to the mother.

### 2.6. Effect size calculation

To quantify the relationship between egg hormones and fitness, we combined both Pearson’s correlation coefficients *r* and biserial correlations *r*bis – throughout we refer to simply *r* for simplicity. First, for correlation coefficients other than Pearson’s *r* (e.g., Spearman’s rho correlation coefficient), percentages, absolute values or inferential statistics (e.g., F-value, t-value, Chi-square value), we calculated Pearson’s *r* using the equations from Nakagawa & Cuthill (2007) and Lajeunesse (2013; Supplementary Information S7). For tests that lacked information on directionality, we determined it from the text, figures, or by directly asking the authors via email. We then calculated the variance in Pearson’s *r* (*Vr*) as V*r* = (1 – *r*^2^) ^2^ ÷ (N – 1) (Borenstein *et al*. 2009). Second, for experimental studies in which two groups were compared (control *vs* treatment), we transformed the means, SDs, and sample sizes of the fitness proxies of interest of each group into biserial correlations coefficients and calculated their sampling variance following Jacobs & Viechtbauer (2017) with the function ‘escalc’ from the R package “metafor” v.2.4-0 (Viechtbauer 2010). Biserial correlations are conceptually equivalent and directly comparable to Pearson’s *r* (Jacobs & Viechtbauer 2017). Meta-analyses involving both Pearson’s and biserial correlation coefficients need to be based on the raw coefficients; this is why we did not use Fisher’s *r*-to-*z* transformation (Jacobs & Viechtbauer 2017). When more than two groups were compared (e.g., control *vs* treatment 1 *vs* treatment 2), we calculated effect sizes between the control and each experimental group and divided the sample size of the control group by the number of times the control group was used (N/2 in the previous example) to account for shared-control non-independence. Whenever more than one effect size could be extracted or calculated for the same data (e.g., when the same result was provided using more than one statistical model), we chose one source using the following order of preference: (a) Pearson’s *r*; (b) other correlation coefficients, (c) mean/percentages/absolute differences between control and experimental groups, and (d) inferential statistics. The sample size assigned to each effect size (i.e., number of clutches or number of offspring) reflected the replication level for the specific trait and experimental design reported by the authors in the papers. If the sample size was not reported at that level or the reported sample size did not correspond to the correct replication level, we used the sample size for the other replication level (if available) or estimated it from the reported degrees of freedom (df + 1) whenever possible (18 effect sizes).

In all cases, effect sizes were coded so that fitness proxies were positively correlated with overall fitness. Specifically, for ’egg mortality,’ ’hatching failure,’ and ’offspring mortality,’ the sign of the effect sizes (k = 5 effect sizes, N = 3 studies) was inverted to ensure comparability across all fitness proxies.

### 2.7. Phylogenies and variance-covariance matrices

We built phylogenetic trees by searching for species in the Open Tree Taxonomy (Rees & Cranston 2017) and extracting their phylogenetic information from the Open Tree of Life (Hinchliff *et al*. 2015) using the R package “rotl” v.3.0.5 (Michonneau *et al*. 2016). We calculated tree branch length following Grafen (1989) with the R package “ape” v.5.4-1 (Paradis & Schliep 2019) and constructed a phylogenetic correlation matrix that was included in all the phylogenetic multilevel models (Figure 1). In addition, we specified sampling variances as a variance-covariance matrix rather than a vector of sampling variances. This variance-covariance matrix had the sampling variance for each effect size on the diagonal and the covariance between these measures, which was calculated assuming a 0.5 correlation between effect size sampling variances from the same study (Noble *et al*. 2017), as off-diagonal elements (see Supplementary Information S8 for a sensitivity analysis using a vector of sampling variances rather than the variance-covariance matrix).

**Figure 1:**
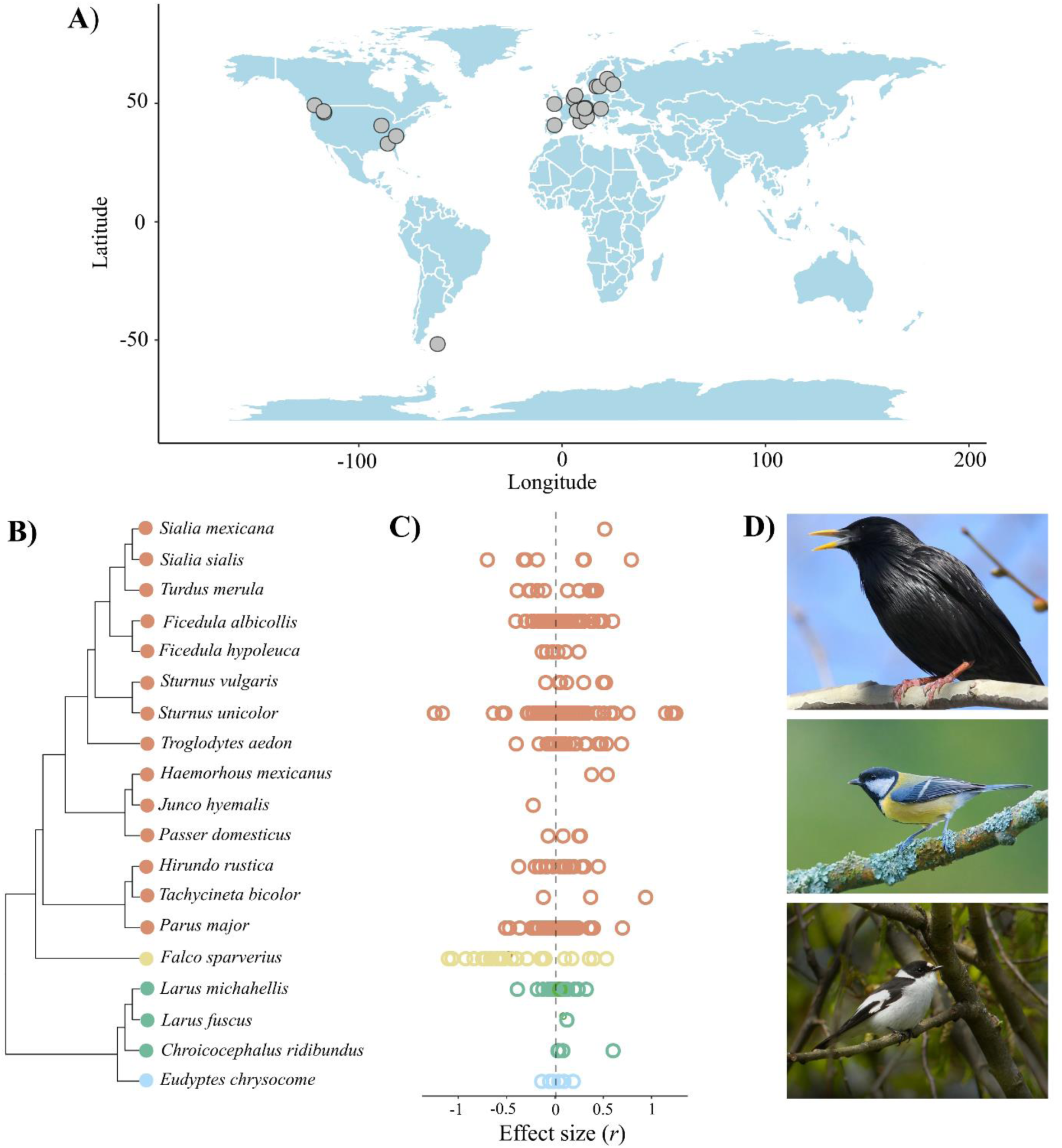
Overview of the meta-analytic data analyzed to test the fitness consequences of maternal egg hormones. A) Global map showing the location of studies included in the meta- analysis that provided longitude and latitude data. Each point represents one study area for which one or more effect sizes were obtained. (B) Phylogenetic tree of the 19 bird species included in the meta-analysis, along with (C) their respective effect sizes. Each color represents a different bird order (red: Passeriformes, yellow: Falconiformes, green: Charadriiformes, and blue: Sphenisciformes). D) Images of the bird species that contributed to most of the data (top to bottom: *Sturnus unicolor*, *Parus major*, *Ficedula albicollis*). All images are copyright-free (CC BY- SA 4.0 DEED. Authors: Peterwchen, Luc Viatour, Frank Vassen, respectively) and were extracted from www.commons.wikipedia.org

### 2.8. General information on analyses

All phylogenetic multilevel meta-analyses and meta-regressions were run using the R package “metafor” v.2.4-0 (Viechtbauer 2010) and included the following random effects: (a) study ID, which encompasses effect sizes extracted from the same study, (b) laboratory ID, which contains effect sizes obtained from the same laboratory, (c) population ID, which considers effect sizes derived from the same population, (d) species name, which encompasses effect sizes derived from the same species, (e) phylogeny, which consists of the phylogenetic correlation matrix and informs about the relatedness between species, and (f) effect ID, which is a unit-level effect and represents residual/within-study variance.

We present our results by first describing the direction and magnitude of the effect, followed by the statistical significance, and finally by reporting the heterogeneity around the mean estimates. This approach places less focus on dichotomizing the results (discussed by e.g., Amrhein *et al*. 2019) and provides crucial insights into the variance across effect sizes, helping assess the generalizability of our findings (discussed by e.g., Gurevitch *et al*. 2018). We provide information on the effect size, both 95% confidence intervals (CI) and 95% prediction intervals (PI), and p-values. Confidence intervals show the range in which the overall average effect is likely to be found, whereas prediction intervals (which incorporate heterogeneity) estimate the likely range in which 95% of effects are expected to occur in studies (past or future) similar to those included in the current study (IntHout *et al*. 2016). For the intercept-only models, we first performed the Cochrane’s *Q* test and then calculated total heterogeneity (σ^2^), its source (or relative heterogeneity; *I*^2^) and two different metrics to understand its magnitude – a mean-standardized metric (*CVH2*) and a variance-mean-standardized metric (*M2*; more in Yang *et al*. 2024). For meta-regression models, we estimated the percentage of heterogeneity explained by the moderators as *R*^2^ (Nakagawa & Schielzeth 2013). We graphically presented the results as orchard plots using the R package “orchaRd” v.2.0 (Nakagawa *et al*. 2023), which visualizes the overall effect size (i.e., meta-analytic mean), along with its 95% confidence and prediction intervals, and individual effect sizes scaled by their precision (i.e., 1/SE).

### 2.9. Main effect model

To estimate the overall effect size for the association between egg hormones and fitness across studies (BH.1), we fitted a phylogenetic multilevel intercept-only meta-analytic model for our entire dataset (k = 438 effect sizes, N = 57 studies). Note that this model was not pre-registered and includes data for all hormones. We used this approach to understand the overall heterogeneity in our data before we proceeded to explain it with our meta-regressions (see below).

To assess the robustness of our results to the choice of effect size, we additionally ran four sensitivity models that, although not stated in our pre-registration, we considered important to confirm that our results were robust to the choice of effect size. For that, we ran four intercept-only models using three subsets of the data: i) Pearson’s *r* and ii) Fisher’s *Zr* (k = 164, N = 39), iii) biserial correlations (k = 272, N = 38), and iv) log-response ratio coefficients (lnRR, k = 269, N = 37; calculated using the function ‘escalc’ from the R package “metafor” v.2.4-0, Viechtbauer 2010). These four sensitivity models reached similar conclusions as the main effect model (Supplementary Information S9).

### 2.10. Biological predictions

To investigate the association between maternal egg hormones and offspring fitness proxies (BP.1), we use data only including effect sizes on offspring fitness (k = 352, N = 42), and then split it into three subsets, one per hormone type: androgens (i.e., androstenedione, 5ɑ- dihydrotestosterone and testosterone; BP.1.1), glucocorticoids (BP.1.2) and thyroids (i.e., TH3 and TH4; BH.1.3). Next, we fitted three multilevel intercept-only meta-analytic models. Note that in our pre-registration we also proposed to study egg progesterone and estradiol (BP.1.4 & BP.1.5); however, we could not because no study fulfilled our eligibility criteria. To further assess the robustness of our results to our choice of analytical strategy for these three hypotheses (BP.1.1 - BP.1.3), we ran a phylogenetic multilevel meta-regression with ‘hormone type’ (levels: androgens, glucocorticoids and thyroids) as a moderator. Although the mean estimate for glucocorticoids (k = 56, N = 8) and thyroids (k = 36, N = 3) was slightly different between the two analytical approaches, likely due to the much smaller sample sizes for those hormones compared to androgens (k = 260, N = 33), the estimates remained statistically non-significant (Supplementary Information S8 & S10).

To test if maternal egg hormones have age-specific fitness consequences for the offspring (BP.3) we fitted two meta-regression models. In the first model (BP.3.1), we originally planned to test if the hormone-fitness relationship differed depending on ‘hormone type’ (levels: androgens, glucocorticoids, and thyroids), ‘offspring life history stage’ (levels: hatching, before independence, and after independence), and their interaction. However, our final model contained only androgen data and the moderator ‘offspring life history stage’ (k = 260, N = 33) because we did not have enough data points for the levels glucocorticoids, thyroids, and after independence (i.e., we had less than 5 data points per level; more in our pre-registration Mentesana *et al*. 2021b). In the second model (BP.3.2), we included ‘hormone type’ (levels: androgens, glucocorticoids and thyroids), ‘offspring relative age’ as a continuous moderator, and their interaction (k = 218, N = 19).

### 2.11. Methodological predictions

We performed three phylogenetic multilevel meta-regressions to investigate the impact of methodological differences on the relationship between maternal egg hormones and both maternal and offspring fitness proxies. In the first model (MP.1), we included ‘hormone type’ (levels: androgens, glucocorticoids and thyroids), ‘study type’ (levels: correlational and experimental), and their interaction as moderators (k = 397, N = 53). In the second model (MH.2), we fitted ‘hormone type’, ‘experimental study object’ (levels: egg and mother), and their interaction as moderators (k = 247, N = 35). Note that due to a lack of correlational studies testing for the effects of thyroids on fitness and of experimental studies manipulating maternal thyroid hormones, we could only test the first two models (MP.1 & MP.2) for androgens and glucocorticoids. In the third model (MP.3), we originally planned to include ‘hormone type’, ‘experimental dose’ (levels: within natural range and supraphysiological), and their interaction as moderators; however, we could only do so for androgens (k = 89, N = 19) due to a lack of data for the other two hormones, and thus, our final model only included ‘experimental dose’ as the only moderator.

### 2.12. Exploratory hypotheses

We performed three additional pre-registered biological and methodological exploratory meta-regressions for which we did not have directional predictions (Mentesana *et al*. 2021b). To test the effect of maternal egg hormones on maternal fitness (BEH.1) we only included effect sizes on maternal fitness proxies (k = 86, N = 46). We originally planned to run a phylogenetic multilevel meta-regression with ‘hormone type’ (levels: androgens, glucocorticoids and thyroids) as a moderator. However, we could only test this for androgens and glucocorticoids (k = 83, N = 44). To test if the effect of maternal egg hormones on offspring fitness differs depending on the matrix in which the hormone was measured/manipulated (BEH.2), we planned to run a phylogenetic multilevel meta-regression using ‘hormone type’, ‘egg site measured’ (levels: albumen and yolk), and their interaction as moderators. Unfortunately, the data set covered information only about glucocorticoids and for a limited number of studies (k = 38, N = 4), preventing us from drawing reliable conclusions (Supplementary Information S11). Finally, to test the effect of maternal egg hormones on fitness depending on the sampling technique used, we ran a phylogenetic multilevel meta-regression using ‘egg sampling method’ (levels: biopsy and entire egg removed) as moderator (MEH.1), and we were able to test this only for androgens (k = 91, N = 9).

### 2.13. Publication bias tests

We tested for evidence of publication bias (following Nakagawa *et al*. 2022; Sánchez-Tójar *et al*. 2018). We did not find any compelling evidence of publication bias either when testing for small study, decline or reporting effects (see Supplementary Information S12 for further details on our approach and the results).

## 3. Results

Overall, we obtained 438 effect sizes from 57 studies across 19 bird species, 4 bird orders, and 34 different populations (Figure 1). All studies but one were conducted in the Northern Hemisphere. Three species accounted for 61% of the effect sizes (i.e., *Sturnus unicolor*, k = 105 effect sizes; *Parus major*, k = 94; *Ficedula albicollis*, k = 69). Androgens were the most studied hormones (k = 333), followed by glucocorticoids (k = 66) and thyroid hormones (k = 39), and most effect sizes (76%) were obtained from experiments (k = 335). The fitness proxies studied corresponded mainly to offspring (k = 352) rather than mothers (k = 86), and those proxies were primarily obtained before offspring independence (k = 223) and by pooling offspring sexes (k = 309).

### 3.1. Main effect model

The intercept-only model revealed a negative, near-zero, and statistically nonsignificant association between egg hormones and fitness - both offspring and maternal - across wild birds (Figure 2; *r* = -0.072, 95% CI: [-0.331, 0.186], 95% PI: [-0.851, 0.707], p-value = 0.583, k = 438, n = 57). Total absolute heterogeneity (σ^2^ = 0.14; *Q* = 48122, p-value < 0.001), relative heterogeneity (*I*^2^ = 95.3%), and the two magnitude metrics (*CVH2* = 26.7, *M2* = 0.96) showed high values of heterogeneity that were beyond the 75^th^ percentile of empirically derived ones in ecological and evolutionary meta-analyses (more in Yang *et al*. 2024; for details see Supplementary Figure 4 & Supplementary Information S13).

**Figure 2:**
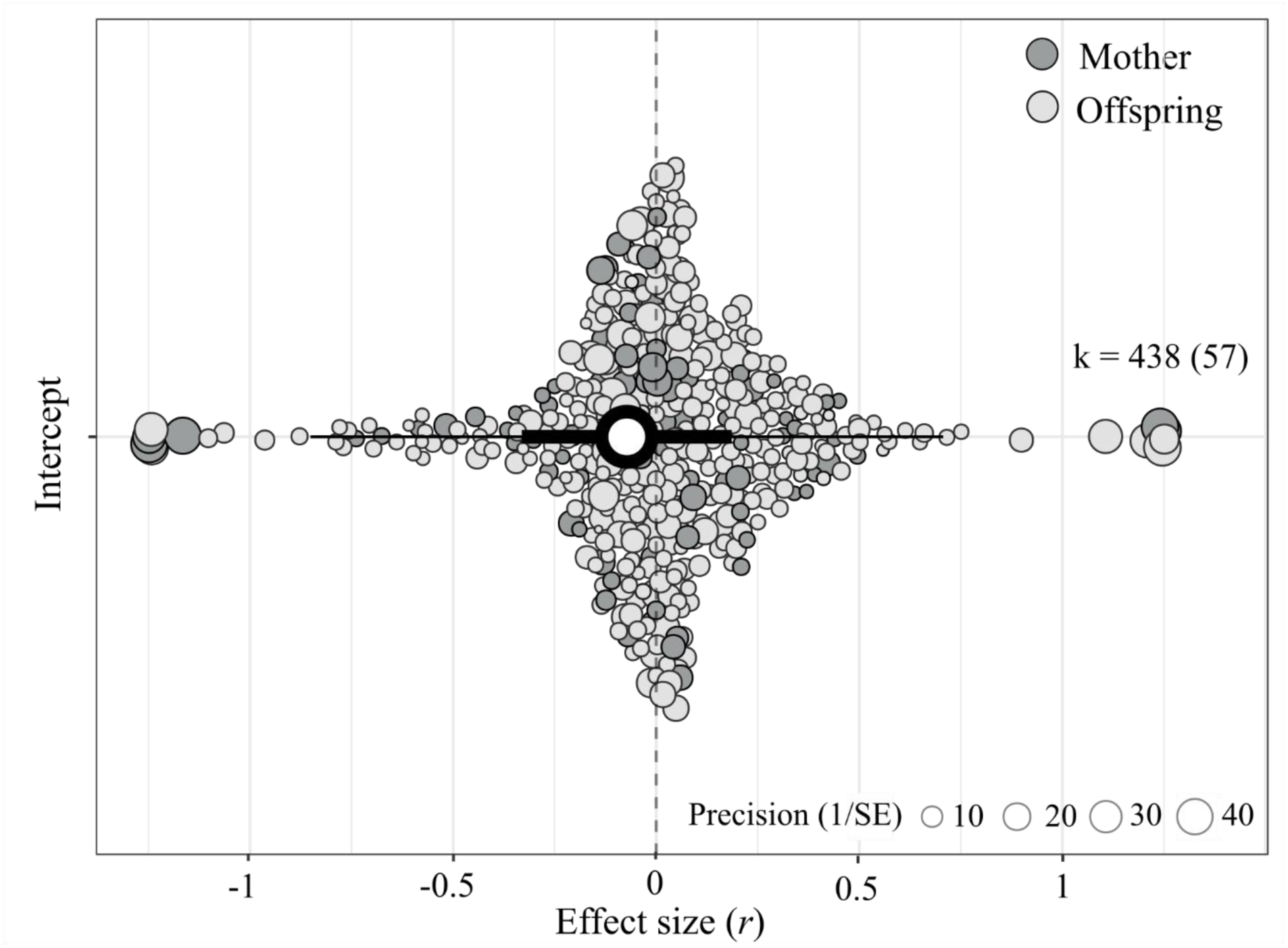
Overall, egg hormone concentrations (androgens, glucocorticoids, and thyroid hormones) show a near-zero association with both maternal (dark gray circles; ca. 20% of effect sizes) and offspring (light gray circles; ca. 80% of effect sizes) fitness in wild birds. Orchard plot of the phylogenetic multilevel (intercept-only) model showing the meta-analytic mean (black circle), 95% confidence intervals (thick whisker), 95% prediction intervals (thin whisker), and individual effect sizes scaled by their precision (gray circles). k is the number of individual effect sizes, the number of studies is shown in brackets.

### 3.2. Biological predictions

#### 3.2.1. Offspring fitness effect sizes (BP.1)

A non-pre-registered intercept-only model confirmed the negative, near-zero, and statistically nonsignificant association between egg hormones and offspring fitness across wild birds (*r* = - 0.063, [95% CI = -0.410, 0.283], [95% PI = -0.928, 0.801], p-value = 0.719, k = 352; N = 42) as well as its high levels of heterogeneity (σ^2^ = 0.16; Q = 22407, p-value < 0.001; *I*^2^total = 95.5%; *CVH2*total = 40.4, *M2*total = 0.9). Moreover, effect sizes for the association between all three types of egg hormones and offspring fitness remained small and statistically nonsignificant (Figure 3; details in Supplementary Information S14) and the moderator hormone type explained negligible heterogeneity (*R*^2^marginal = 0.3%). Moreover, none of the three estimates differed statistically from each other (p-value androgens *vs* glucocorticoids = 0.400; androgens *vs* thyroids = 0.533; glucocorticoids *vs* thyroids = 0.970).

**Figure 3:**
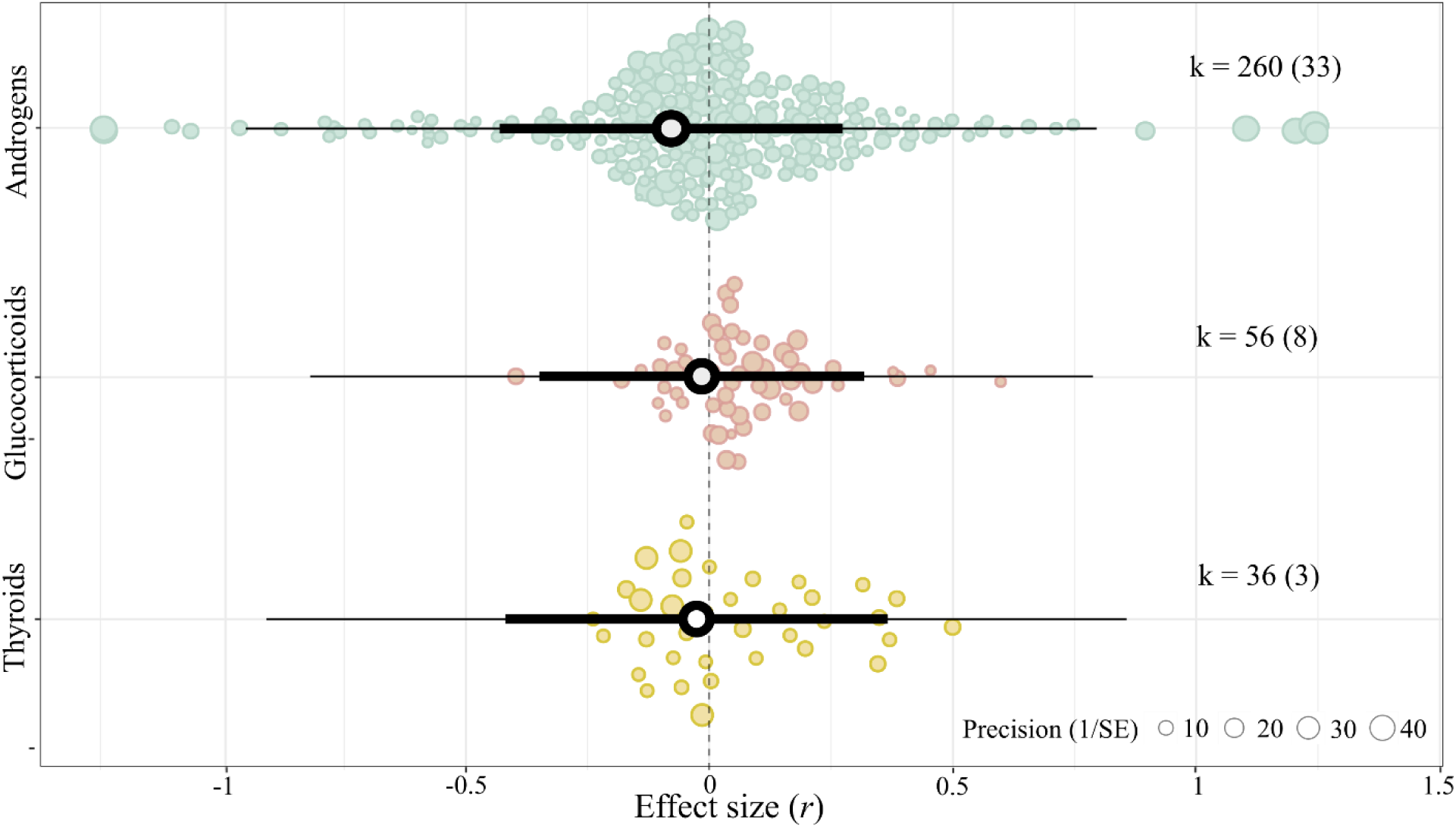
Regardless of hormone type, egg hormone concentrations show a near-zero association with offspring fitness in wild birds. Orchard plot of the phylogenetic multilevel meta-regression showing the mean estimates (black circles), 95% confidence intervals (thick whisker), 95% prediction intervals (thin whisker), and individual effect sizes scaled by their precision (coloured circles). k is the number of individual effect sizes, the number of studies is shown in brackets.

#### 3.2.3. Age-specific offspring fitness effect sizes (BP.3)

For both altricial and precocial species, androgens (the only hormone for which we had enough data at different categorical age stages to perform this test; BP.3.1) seemed to be positively associated with offspring fitness at hatching and negatively associated before and after independence. However, effect sizes were small and statistically nonsignificant (Figure 4A; hatching: *r* = 0.009, [95% CI = -0.389, 0.407], [95% PI = -0.937, 0.956], p-value = 0.963, k = 21, N = 7; before independence: *r* = -0.076, [95% CI = -0.446, 0.294], [95% PI = -1.012, 0.859], p-value = 0.684, k = 223, N = 30; after independence: *r* = -0.174, [95% CI = -0.618, 0.271], [95% PI = -1.141, 0.793], p-value = 0.442, k = 16, N = 4), and the moderator explained negligible heterogeneity (*R*^2^marginal = 0.6%). None of the three age stages differed statistically from each other (p-value hatching *vs* before independence = 0.334; hatching *vs* after independence = 0.240; before *vs* after independence = 0.448).

**Figure 4:**
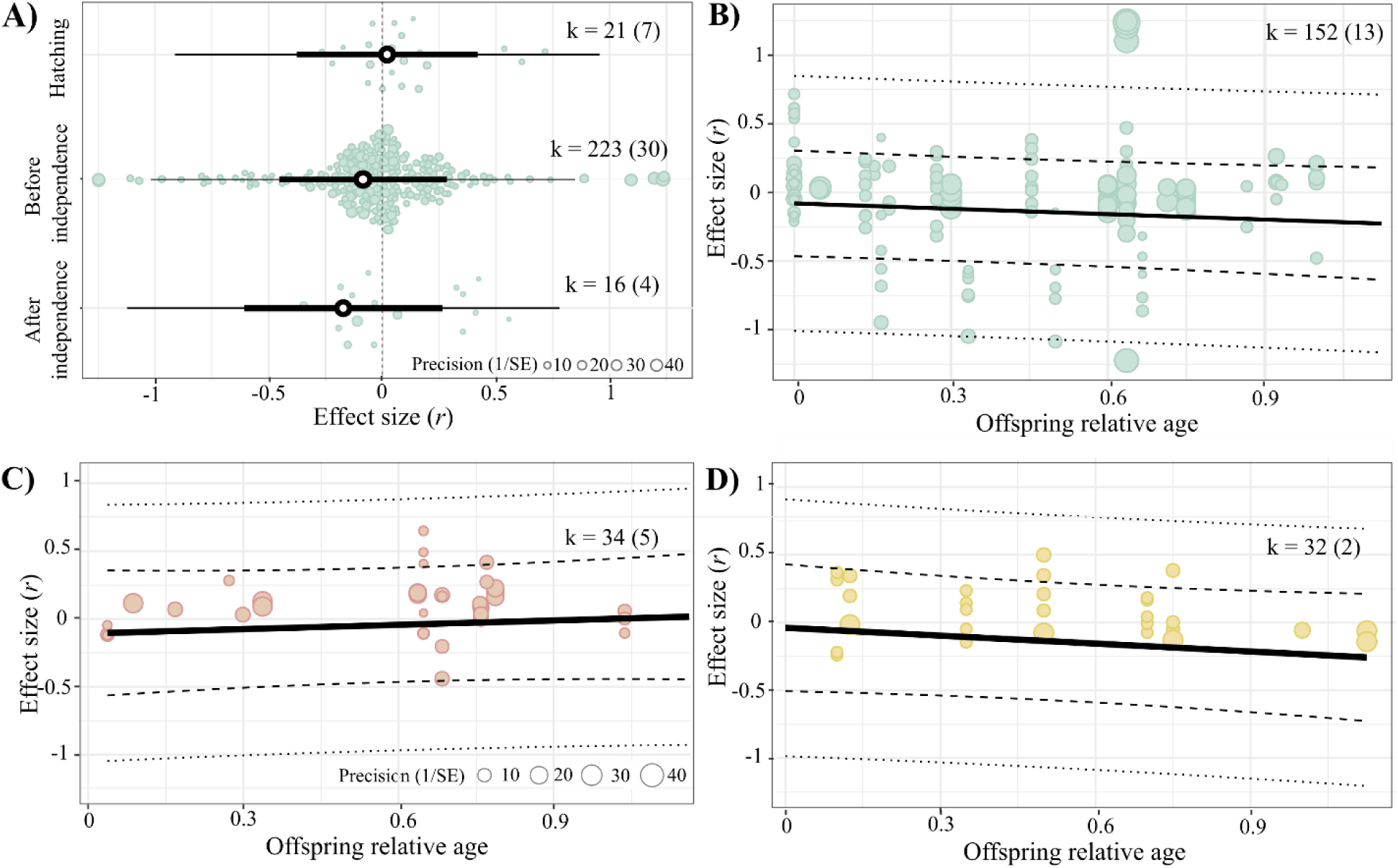
Regardless of offspring age and hormone type, egg hormone concentrations show a near-zero association with offspring fitness in wild birds. A) Egg androgen concentrations and offspring fitness in both altricial and precocial species. Orchard plot of the phylogenetic multilevel meta-regression displaying mean estimates (black circles), 95% confidence intervals (thick whiskers), 95% prediction intervals (thin whiskers), and individual effect sizes scaled by precision (colored circles). Panels B-D refer to B) androgen, C) glucocorticoid and D) thyroid egg hormone concentrations with offspring fitness in altricial species. We plotted a separate bubble plot for each of the hormone types analyzed in a single phylogenetic multilevel meta-regression, showing model estimates (solid lines), 95% CIs (thick dashed lines), 95% prediction intervals (thin dashed lines), and individual effect sizes by precision (colored circles: green for androgens, red for glucocorticoids, yellow for thyroids). k is the number of individual effect sizes, the number of studies is shown in brackets.

In altricial species, the association between egg androgen or thyroid concentrations and offspring fitness proxies appeared to weaken throughout the nesting phase, while the association with glucocorticoids showed the opposite trend (BP.3.2); however, none of those slopes were statistically significant (Figure 4B-D; Table 2) and the moderator only explained little heterogeneity (*R*^2^ = 1.7%).

**Table 2:** Results of phylogenetic multilevel meta-regression models testing whether the association between egg hormone concentrations and offspring fitness vary throughout the nesting phase in altricial bird species^‡^.

|  | Estimate | SE | t-value | df | p-value | L 95% CI | U 95% CI |
| --- | --- | --- | --- | --- | --- | --- | --- |
| Intercept (Androgens) | -0.088 | 0.197 | -0.445 | 212 | 0.657 | -0.476 | 0.301 |
| Glucocorticoids | -0.005 | 0.152 | 0.033 | 212 | 0.974 | -0.295 | 0.305 |
| Thyroids | 0.071 | 0.153 | 0.465 | 212 | 0.643 | -0.231 | 0.374 |
| Offspring age | -0.132 | 0.099 | -1.331 | 212 | 0.185 | -0.327 | 0.063 |
| Glucocorticoids * Offspring age | 0.224 | 0.220 | 1.019 | 212 | 0.309 | -0.209 | 0.658 |
| Thyroids * Offspring age | -0.060 | 0.209 | -0.289 | 212 | 0.773 | -0.473 | 0.352 |

<sup>‡</sup> df: degrees of freedom; CI: confidence interval. Offspring age was standardized as the age of the offspring divided by the average nesting time and was calculated for each study based on information reported by the authors (more in Table S5.1).

### 3.3. Methodological predictions

#### 3.3.1. Fitness effect sizes in correlational *vs* experimental studies (MP.1)

For the two hormone types for which sufficient data existed, androgen and glucocorticoids, the associations between concentrations and both maternal and offspring fitness seemed negative, near-zero, and statistically nonsignificant (Figure 5A; Supplementary Information S15A). The moderator explained little heterogeneity (*R*^2^ = 1.4%), and the differences between correlational and experimental were statistically nonsignificant regardless of hormone type (p-value androgens = 0.142; glucocorticoids = 0.340).

**Figure 5:**
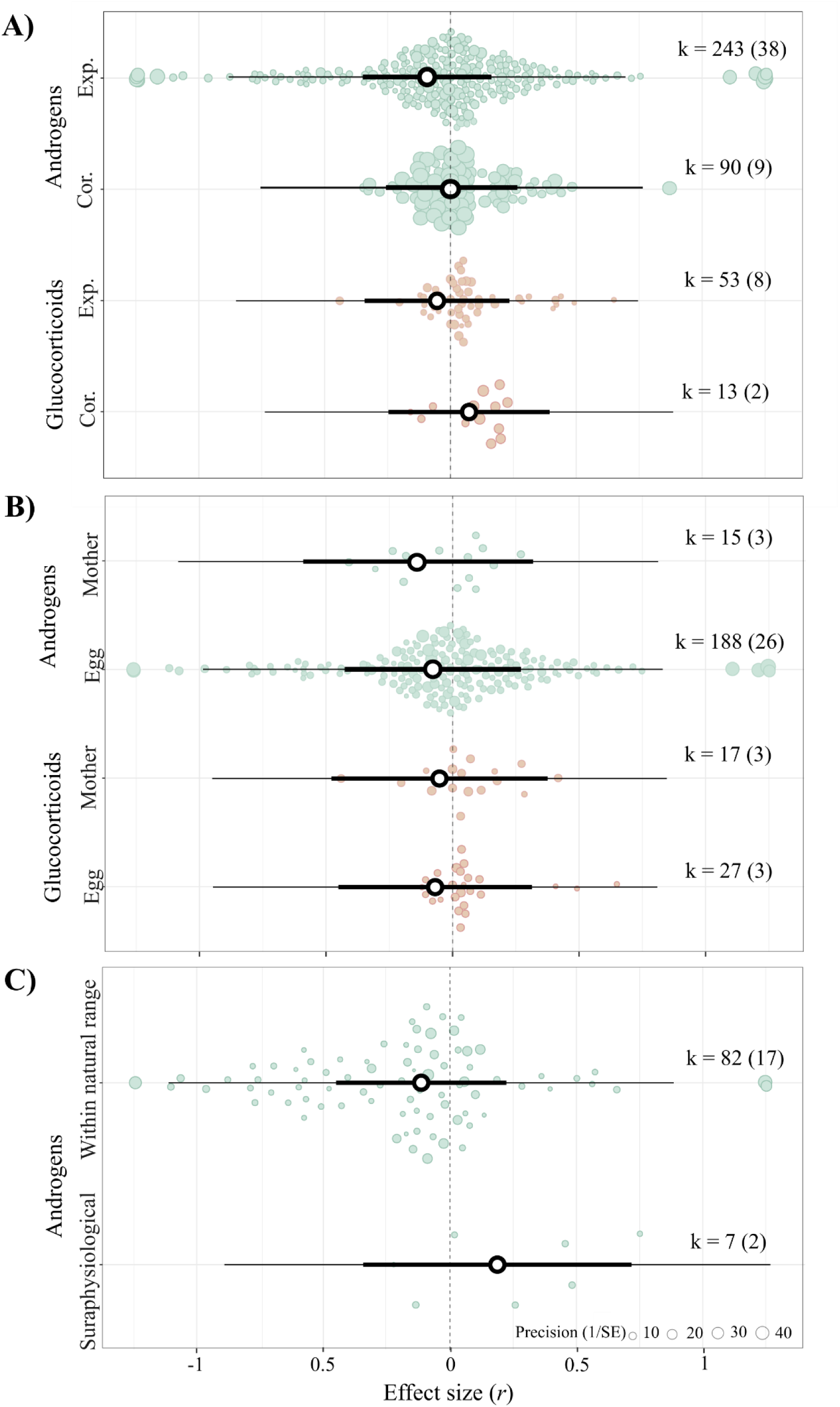
The relationship between androgen and glucocorticoid egg hormone concentrations and fitness is near-zero, irrespective of A) the type of study performed (i.e., correlational or experimental), B) the study object on which the experiment was performed (i.e., the mother or the egg), and, solely for androgens, C) the experimental dose used (i.e., within or above the natural range). Orchard plots of three phylogenetic multilevel meta-regression models showing mean estimates (black circles), 95% confidence intervals (thick whisker), 95% prediction intervals (thin whisker), and individual effect sizes scaled by their precision (coloured circles). k is the number of individual effect sizes, the number of studies is shown in brackets.

#### 3.3.2. Fitness effect sizes in experimental studies performed on the mother *vs* the egg (MP.2)

The negative but statistically nonsignificant association between egg androgen or glucocorticoid concentrations and offspring fitness remained independent of whether hormonal manipulations were performed on the mother or the egg (Figure 5B; Supplementary Information S15B; p-value mother *vs* eggs = androgens 0.654, glucocorticoids 0.985). The moderator explained negligible heterogeneity (*R*^2^ = 0.2%).

3.3.3. Fitness effect sizes in experimental studies conducted within *vs* above the natural physiological range (MP.3)

Although the association between egg androgens and offspring fitness appeared to be positive at doses above the physiological range and seemed negative at doses within the natural range, none of the estimates were statistically significant (Figure 5C; Supplementary Information S15C) or differed from each other (p-value = 0.175). Moreover, the moderator explained little heterogeneity (*R*^2^marginal = 2.9%).

### 3.4. Exploratory biological and methodological hypotheses

The association between egg hormones and maternal fitness was statistically nonsignificant both for androgens and glucocorticoids (BEH.1; androgens: *r* = -0.052, [95% CI = -0.159, 0.054], [95% PI = -0.826, 0.722], p-value = 0.332, k = 73, N = 36; glucocorticoids: *r* = 0.082, [95% CI = -0.183, 0.348], [95% PI = -0.729, 0.893], p-value = 0.540, k = 10, N = 8). The moderator explained negligible heterogeneity (*R*^2^ = 1.3%), and the two hormone groups did not differ statistically from each other (p-value = 0.351).

We found that the association between maternal egg androgen concentrations on fitness did not differ statistically between the two sampling techniques used (p-value = 0.196; MEH.1). When androgens were measured using an egg biopsy, they seemed to be negatively associated with fitness, whereas when the entire egg was removed, they appeared to be positively associated with fitness (egg biopsy: *r* = -0.202, [95% CI = -0.681, 0.278], [95% PI = -1.025, 0.622], p-value = 0.406, k = 19, N = 2; entire egg removed: *r* = 0.144, [95% CI = -0.089, 0.377], [95% PI = -0.565, 0.853], p-value = 0.223, k = 74, N = 8; *R*^2^ = 14.7%).

Due to small sample sizes, these results should be interpreted cautiously as the models likely are heavily underpowered.

## 4. Discussion

Our pre-registered systematic review and meta-analysis found that the fitness effects of egg hormones across 19 wild bird species are neither as strong nor as clear as previously suggested. The overall mean effect of androgens, glucocorticoids, and thyroid hormones on both maternal and offspring fitness was statistically indistinguishable from zero. However, those effects were also highly heterogeneous, suggesting that the magnitude and direction of the relationship between hormone concentrations and fitness is context specific. We tested several biological and methodological moderators that we predicted could explain the differences in fitness effects for offspring, for which most of the data exists, but they only accounted for a small percentage of the heterogeneity. Below, we discuss our findings in detail, examining our hypotheses in specific contexts, and highlighting knowledge gaps, biases in the current literature, and areas for future research.

The influence of egg hormone concentrations on fitness has historically been expected to vary by hormone group: increased egg androgens and thyroid hormones are predicted to enhance fitness, while increased glucocorticoids are expected to reduce it (e.g., von Engelhardt & Groothuis 2011; Groothuis *et al*. 2005a; Haussmann *et al*. 2012; Hsu *et al*. 2017; Polich *et al*. 2018; Ruuskanen *et al*. 2018; Stier *et al*. 2009). However, our results showed that the relationship between hormone concentration and offspring fitness proxies was similarly small and statistically indistinguishable from zero for all three hormone groups, with ’hormone type’ explaining a negligible 0.4% of the heterogeneity among effect sizes. Our exploratory analysis testing the relationship with maternal fitness for the two hormones for which sufficient data were available (i.e., androgens and glucocorticoids) also showed a small and statistically nonsignificant relationship. In all, our results, including all sensitivity analyses, robustly show no support for a consistent overall relationship between egg hormones and fitness in wild bird populations.

Egg hormone concentrations are also suggested to have sex-specific effects on offspring fitness (reviewed by e.g., von Engelhardt & Groothuis, 2011). While we aimed to test this hypothesis (see BP.2 in the Introduction), data on offspring sex-specific effects were available in only 12.2% of cases (43 out of 352 offspring effect sizes, 7 studies). This limited dataset included only two hormone types (androgens and thyroid hormones) and three bird species (*Parus major*, *Ficedula albicollis*, and *Falco sparverius*). The relative availability of sex-specific data in our study was even lower than the 32% reported by Podmokła et al. (2018) for experimental studies conducted primarily in captive birds. Notably, while Podmokła et al. (2018) included data on androgens and glucocorticoids, we found information only for androgens and thyroid hormones, and overall, 56.1% of the studies in our dataset were not included in theirs. Altogether, this clearly indicates the current lack of sex-specific data for all egg hormones in both captive and wild birds, which limits our understanding of whether maternal egg hormones have sex-specific fitness consequences.

The relationship between egg hormones and offspring fitness proxies showed no overall change with age. Theory predicts that maternal effects should weaken as offspring encounter more environments throughout ontogeny (e.g., Mousseau & Dingle 1991), and a meta-analysis of maternal (genetic) effects on vertebrate species supported this prediction (Moore *et al*. 2019). The absence of an ontogenetic decline in our study might be because egg hormones already had a weak effect at hatching, despite being expected to be strongest at this stage compared to later ones. Moreover, Moore *et al*. (2019) found that maternal (genetic) effects declined more for traits with higher maternal influence such as morphological traits compared to other traits such as behavior and physiology. Most studies providing offspring life-stage or age focused on androgens and, thus, we urge researchers to gather more data on glucocorticoids and thyroid hormones along developmental stages to better understand the relationship between hormones and fitness across life stages.

In wild bird populations, the overall null effect between hormones and fitness proxies appears consistent, even when accounting for methodological differences across studies. Specifically, we found that egg androgens and glucocorticoids had no overall effect on fitness proxies regardless of study type (correlational or experimental) or experimental focus (mother or egg). The experimental dose being within the natural range or above physiological concentrations, and the sampling technique used (egg biopsy or entire egg collection) explained little to negligible amounts of the observed heterogeneity. These results are also relevant, as they suggest that manipulating egg hormone concentrations neither leads to fitness gains, as expected when females face physiological constraints or trade-offs, nor incurs fitness costs, as predicted if maternal egg hormones are adaptive (von Engelhardt & Groothuis 2011). A methodological issue that requires further consideration is the variation in extraction methods across laboratories, particularly for glucocorticoids. In immunoassays, glucocorticoid antibodies can cross-react with progesterone and its precursors, leading to an overestimation of yolk glucocorticoid concentrations (Rettenbacher *et al*. 2009, 2013). Column chromatography is recommended to prevent this, but likely due to the complexity of this method, only 2 of 10 laboratories studying egg glucocorticoids and fitness proxies employed it (Bowers et al. 2016; Mentesana et al. 2021a). Overestimated glucocorticoid concentrations resulting from methodological issues might contribute to discrepancies across labs in the observed relationships between egg glucocorticoids and fitness proxies. Supporting this idea, we found that relative heterogeneity was highly explained by differences among laboratories (*I^2^*among-laboratory = 42.17%; Supplementary Information Table S10.2).

By using a pluralistic approach to estimate total heterogeneity and stratify it across the different hierarchical levels explored, we found that the overall association between egg hormones and both maternal and offspring fitness was highly heterogeneous and stemming from within-study variation and phylogenetic relationships among species (Supplementary Information S10.2 & S13). The high levels of heterogeneity associated with within- compared to among-study variation, the latter being zero in all models (and robust to sensitivity tests), suggest that the small and statistically nonsignificant overall effect found by our meta-analysis (i.e., our meta-analytic mean) would be generalizable among studies as long as (i) we define replication as the testing of the null hypothesis at the among-study level and, (ii) importantly, if within-study level variation - both biological and methodological, are accounted for (Yang *et al*. 2024). Our moderator analyses testing for both biological and methodological differences, however, did not explain a substantial amount of heterogeneity, limiting our potential for generalization. We hypothesized that such differences may stem from researchers reporting multiple fitness proxies within studies (mean ± SD = 2.96 ± 1.86) and conducted an additional (non-pre-registered) meta-regression using the entire dataset. None of the fitness proxy-specific mean estimates were statistically significantly different from zero (see Supplementary Figure 5; Supplementary Information S16), and the fitness proxy used accounted for a small but somewhat non-negligible percentage of the heterogeneity (*R*²marginal = 3.5%). Among-species differences, and particularly, phylogenetic relationships among the 19 bird species were associated with high levels of heterogeneity, which indicates that the size and magnitude of the overall association between egg hormones and fitness proxies differ among taxonomic groups (Figure 1C). We explored whether species-specific characteristics that are strongly correlated with phylogeny may explain part of that heterogeneity. For this, we ran a (non-pre-registered) model to compare precocial *vs* altricial species and found that developmental mode explains little heterogeneity (*R*²marginal = 0.3%; see Supplementary Information S17). To better understand the factors driving these differences, future research could focus on understanding if species-specific characteristics (e.g., life-history, ecology) might be driving the phylogenetic signal observed in our study as well as investigate whether egg hormones and fitness are associated in other bird species not yet studied.

Three additional factors might explain the heterogeneity that we found across effect sizes. First, one complex factor that we could not account for is the environment experienced by mothers and offspring. Theory predicts that anticipatory maternal effects evolve when mothers and offspring encounter similar environments, or when environmental fluctuations are low (Hoyle & Ezard 2012; Kuijper & Hoyle 2015). Egg hormones may significantly influence fitness proxies only when maternal and offspring environments match, and/or when mothers can predict the environment their offspring will encounter. Conversely, maternal effects may be non-anticipatory, with heterogeneity in fitness outcomes arising from the cost-benefit balance of egg hormones under varying environmental scenarios experienced solely by the offspring (reviewed by Groothuis *et al*. 2020). Second, maternal effects are multivariate. Mothers influence their offspring’s prenatal environment by depositing various compounds into the egg besides hormones (e.g., immunoglobulins, antioxidants, fatty acids, proteins; Lovern & Wade 2001; McCormick 1999; Mentesana *et al*. 2019; Valcu *et al*. 2019; Weiss *et al*. 2011), which may interact (Groothuis *et al*. 2019; Mentesana *et al*. 2021a; Williams & Groothuis 2015). If these egg components do not act independently on fitness traits, and their correlations are weak or different combinations have similar outcomes, testing the effects of single components independently, as we did, is likely to result in high heterogeneity. Third, there is growing evidence that embryos are not passive responders to maternal signals (Bebbington & Groothuis 2021; Groothuis *et al*. 2019). Embryos seem adapted to receiving maternal hormone signals, as they have hormone receptors before producing their own hormones (Godsave *et al*. 2002; Kumar *et al*. 2019). However, recent evidence suggests that embryos can rapidly metabolize maternal hormones in the egg into potentially inactive steroid forms (Kumar *et al*. 2018; Vassallo *et al*. 2014). If the hormone concentrations in the egg do not reflect the amount actually incorporated by the offspring, this could explain the variation in effect sizes and the overall lack of effect that manipulating egg hormone concentrations has on offspring fitness.

Egg hormone-mediated maternal effects are considered an adaptive mechanism, where mothers adjust hormone concentrations in response to environmental cues to optimize maternal and/or offspring fitness. This assumes that mothers can modify hormone levels based on the environment, in addition to the maternal effects on fitness explored in this study. However, it is debated whether hormone deposition is truly female-controlled (von Engelhardt & Groothuis 2011; Gil 2008; Groothuis & Schwabl 2008). Groothuis & Schwabl (2008) proposed three mechanisms: (1) egg hormone concentrations mirror maternal plasma levels, (2) eggs act as a sink for excess steroids, or (3) egg and maternal hormone levels are independently regulated. The first two mechanisms suggest egg hormones reflect maternal responses to the environment or physiology, not being offspring-specific. If mothers cannot adjust hormone transfer, adaptive maternal effects may be constrained. Understanding which mechanism is operating and whether it is common among species is pivotal to the study of the correlations between egg hormones concentrations and fitness.

## 5. Conclusions

Thirty years after the discovery of egg hormones, our meta-analysis showed that an increase in their concentrations has a near zero and highly variable overall mean effect on fitness proxies in wild bird populations. We could not identify any predictor explaining much of the high levels of heterogeneity found among effect sizes, even after accounting for various biological (e.g., hormone type and age) and methodological (e.g., study type, experimental subject, dose) factors.

We hope our work helps advance the field of maternal effects via egg hormone deposition and inspires new research into its many intriguing aspects. Key areas for future research include studying phylogenetically distant species, investigating whether the near-zero mean effect of egg hormones on both maternal and offspring fitness proxies can be explained by environmental context, studying sex-specific effects of egg hormones, and clarifying the mechanisms of maternal transfer to eggs. Such research questions are essential for understanding the adaptive significance of maternal effects via egg deposition, so we urge researchers in the field to address them.

## Acknowledgements

We thank all the authors who replied to our email, addressed our questions, and kindly shared their data: Alexandra Bentz, Amber Rice, Barbara Tschirren, Renee Duckworth, Bin-Yan Hsu, E. Diego Rubolini, Keith Bowers, Gergely Hegyi, Ismael Galvan, Jaime Muriel, János Török, Jose Carlos Noguera, Keith W. Sockman, Kristen J. Navara, Marco Parolini, Scott Sakaluk, and Suvi Ruuskanen. We also want to thank Yefeng Yang for his feedback on interpreting heterogeneity results. This research was funded by the Max Planck Gesellschaft (to MH).

## Author’s contributions

Conceptualization: LM, MH and AST. Data curation: LM and AST. Formal analysis: LM and AST. Funding acquisition: MH. Investigation: LM, PBD, and NMA. Methodology: LM and AST. Project administration: LM. Software: LM and AST. Supervision: LM and AST. Validation: LM, PBD, NMA, and AST. Visualization: LM and NMA. Writing – original draft: LM. Writing – review & editing: all authors.

## Data availability statement

All data and code are available on GitHub (https://github.com/ASanchez-Tojar/meta-analysis_egg_hormones_and_fitness). Upon acceptance of this work, the repository will be assigned with a DOI via Zenodo.

## Notes

https://github.com/ASanchez-Tojar/meta-analysis_egg_hormones_and_fitness

### Competing Interest Statement

The authors have declared no competing interest.

### Summary of Updates

In Table 1 we have added information and cites on all the 57 studies used in our meta-analysis; We have moved the analysis about sex-specific effects to Supplementary files because of low statistical power to have confident results; Conclusions remain the same, but the number of effect sizes is different than the previous version because we added data from one study and removed data from another one.

