## Supplementary Information & Figures for "Do egg hormones have fitness consequences in wild birds? A systematic review and meta-analysis"

* Author for correspondence: Dr. Lucia Mentesana

**Supplementary Information S1.** Sex-specific tests of the effects of maternal egg hormones on offspring fitness and results (BP.2).

Sex-specific tests

To test if maternal egg hormones have sex-specific fitness consequences for the offspring (BP.2) we initially planned to run a phylogenetic multilevel meta-regression including 'hormone type' (i.e., androgens, glucocorticoids, thyroids), 'sex', and their interaction as moderators (see pre-registration Mentesana et al. 2021). In addition, we planned to include six random effects (i.e, study ID, laboratory ID, population ID, species name, phylogeny, and effect ID). However, the number of studies for which offspring sex was available is limited (k = 43, N = 7). This small data set covered only two hormones (i.e., androgens and tyroid hormones) and three species (*Parus major*, *Ficedula albicollis*, and *Falco sparverious*). For thyroids specifically, the dataset consisted on only two studies (k = 26 effect sizes) on two species (*P.major* and *F.albicollis*). We therefore had to reduce our model to a multilevel meta-regression including as moderators 'hormone type' (levels: androgen and thyroids), 'sex' (levels: female, male), and their interaction, and only three random effects (i.e., study ID, species name, and effect ID). Given the limited data available, the results of this meta-regression should be interpreted with extreme caution, which is the main reason for why these results are presented in the supplementary material despite BP.2. being part of our original list of pre-registered predictions.

Sex-specific results

The limited data available to test the association between both egg androgens and thyroids with offspring fitness showed that the effect tended to be negative regardless of offspring sex, but none of the estimates were statistically significant (female and male p-values: androgens 0.421 and 0.133, thyroids 0.433 and 0.691). Both moderators combined explained a moderate amount of heterogeneity for this subset of the data ($R^2$marginal = 5.2%). The post-hoc tests showed no statistical difference between males and females for both androgens (p-value = 0.058) and thyroids (p-value = 0.053).

Notice that since the results from the model described above containing interaction terms are not straightforward, below we present the results from a meta-regression model that used a single artificial variable that combined hormone type with sex as the moderator (levels: androgens male, androgens female, thyroid male, and thyroid female).

| | | Mean | L 95% CI | U 95% CI | L 95% PI | U 95% PI | p-value | k | N |
|---|---|---|---|---|---|---|---|---|---|
| Androgens | female | -0.288 | -1.006 | 0.430 | -1.604 | 1.027 | 0.421 | 9 | 5 |
| | male | -0.530 | -1.230 | 0.169 | -1.836 | 0.775 | 0.058 | 8 | 5 |
| Thyroids | female | -0.291 | -1.032 | 0.451 | -1.619 | 1.038 | 0.991 | 13 | 2 |
| | male | -0.147 | -0.888 | 0.595 | -1.475 | 1.182 | 0.471 | 13 | 2 |

† CI: confidence interval; PI: prediction interval; L: lower; U: upper; k = number of effect sizes; N = number of studies.

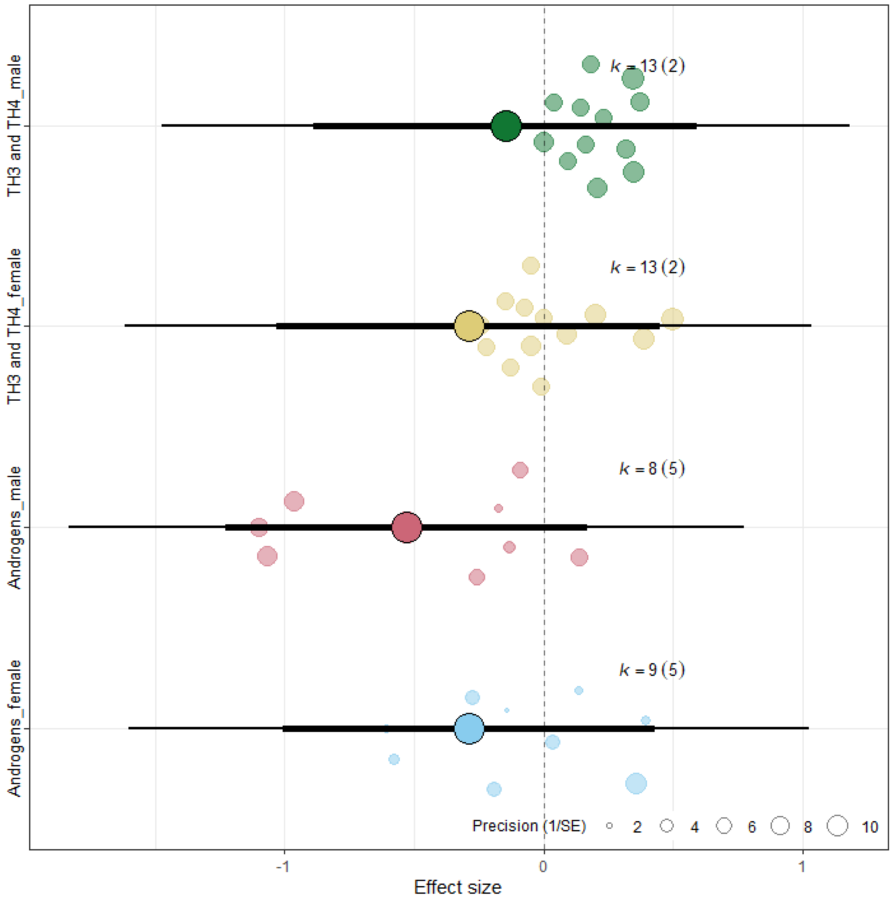


Figure S1: Regardless of offspring sex, androgen and thyroid egg hormone concentrations show a near-zero association with offspring fitness in wild birds. Orchard plots of the phylogenetic multilevel meta-regression showing male and female mean estimates (black circles), 95% confidence intervals (thick whisker), 95% prediction intervals (thin whisker), and individual effect sizes scaled by their precision (coloured circles). k is the number of individual effect sizes, the number of studies is shown in brackets.

**Supplementary Information S2.** Quality assessment of our systematic review and meta-analysis testing the extent to which prenatal maternal hormone deposition into eggs relates to fitness in wild birds. For it, we filled out the ‘Interactive PRISMA-EcoEvo Checklist’ (https://prisma-ecoevo.shinyapps.io/checklist; O’Dea *et al.* 2021).

| Checklist item | Item score | Sub-item number | Sub-item | Reported by authors? |
|---|---|---|---|---|
| Title and abstract | 100% | 1.1 | Identify the review as a systematic review, meta-analysis, or both | Yes |
| | | 1.2 | Summarise the aims and scope of the review | Yes |
| | | 1.3 | Describe the data set | Yes |
| | | 1.4 | State the results of the primary outcome | Yes |
| | | 1.5 | State conclusions | Yes |
| | | 1.6 | State limitations | Yes |
| Aims and questions | 100% | 2.1 | Provide a rationale for the review | Yes |
| | | 2.2 | Reference any previous reviews or meta-analyses on the topic | Yes |
| | | 2.3 | State the aims and scope of the review (including its generality) | Yes |
| | | 2.4 | State the primary questions the review addresses (e.g. which moderators were tested) | Yes |
| | | 2.5 | Describe whether effect sizes were derived from experimental and/or observational comparisons | Yes |

| | | | | |
|---|---|---|---|---|
| Review registration | 100% | 3.1 | Register review aims, hypotheses (if applicable), and methods in a time-stamped and publicly accessible archive and provide a link to the registration in the methods section of the manuscript. Ideally registration occurs before the search, but it can be done at any stage before data analysis. | Yes |
| | | 3.2 | Describe deviations from the registered aims and methods | Yes |
| | | 3.3 | Justify deviations from the registered aims and methods | Yes |
| Eligibility criteria | 100% | 4.1 | Report the specific criteria used for including or excluding studies when screening titles and/or abstracts, and full texts, according to the aims of the systematic review (e.g. study design, taxa, data availability) | Yes |
| | | 4.2 | Justify criteria, if necessary (i.e. not obvious from aims and scope) | Yes |
| Finding studies | 100% | 5.1 | Define the type of search (e.g. comprehensive search, representative sample) | Yes |
| | | 5.2 | State what sources of information were sought (e.g. published and unpublished studies, personal communications) | Yes |
| | | 5.3 | Include, for each database searched, the exact search strings used, with keyword combinations and Boolean operators | Yes |

| | | | | |
|---|---|---|---|---|
| | | 5.4 | Provide enough information to repeat the equivalent search (if possible), including the timespan covered (start and end dates) | Yes |
| Study selection | 100% | 6.1 | Describe how studies were selected for inclusion at each stage of the screening process (e.g. use of decision trees, screening software) | Yes |
| | | 6.2 | Report the number of people involved and how they contributed (e.g. independent parallel screening) | Yes |
| Data collection process | 100% | 7.1 | Describe where in the reports data were collected from (e.g. text or figures) | Yes |
| | | 7.2 | Describe how data were collected (e.g. software used to digitize figures, external data sources) | Yes |
| | | 7.3 | Describe moderator variables that were constructed from collected data (e.g. number of generations calculated from years and average generation time) | Yes |
| | | 7.4 | Report how missing or ambiguous information was dealt with during data collection (e.g. authors of original studies were contacted for missing descriptive statistics, and/or effect sizes were calculated from test statistics) | Yes |
| | | 7.5 | Report who collected data | Yes |

| | | | | |
|---|---|---|---|---|
| | | 7.6 | State the number of extractions that were checked for accuracy by co-authors | Yes |
| Data items | 100% | 8.1 | Describe the key data sought from each study | Yes |
| | | 8.2 | Describe items that do not appear in the main results, or which could not be extracted due to insufficient information | Yes |
| | | 8.3 | Describe main assumptions or simplifications that were made (e.g. categorising both 'length' and 'mass' as 'morphology') | NA |
| | | 8.4 | Describe the type of replication unit (e.g. individuals, broods, study sites) | Yes |
| Assessment of individual study quality | 100% | 9.1 | Describe whether the quality of studies included in the systematic review or meta-analysis was assessed (e.g. blinded data collection, reporting quality, experimental versus observational) | Yes |
| | | 9.2 | Describe how information about study quality was incorporated into analyses (e.g. meta-regression and/or sensitivity analysis) | Yes |
| Effect size measures | 100% | 10.1 | Describe effect size(s) used | Yes |
| | | 10.2 | Provide a reference to the equation of each calculated effect size (e.g. standardised mean difference, log response ratio) and (if applicable) its sampling variance | Yes |

| | | | | |
|---|---|---|---|---|
| | | 10.3 | If no reference exists, derive the equations for each effect size and state the assumed sampling distribution(s) | NA |
| Missing data | 100% | 11.1 | Describe any steps taken to deal with missing data during analysis (e.g. imputation, complete case, subset analysis) | Yes |
| | | 11.2 | Justify the decisions made to deal with missing data | Yes |
| Meta-analytic model description | 100% | 12.1 | Describe the models used for synthesis of effect sizes | Yes |
| | | 12.2 | The most common approach in ecology and evolution will be a random-effects model, often with a hierarchical/multilevel structure. If other types of models are chosen (e.g. common/fixed effects model, unweighted model), provide justification for this choice | NA |
| Software | 80% | 13.1 | Describe the statistical platform used for inference (e.g. R) | Yes |
| | | 13.2 | Describe the packages used to run models | Yes |
| | | 13.3 | Describe the functions used to run models | Yes |
| | | 13.4 | Describe any arguments that differed from the default settings | No |
| | | 13.5 | Describe the version numbers of all software used | Yes |

| | | | | |
|---|---|---|---|---|
| Non-independence | 100% | 14.1 | Describe the types of non-independence encountered (e.g. phylogenetic, spatial, multiple measurements over time) | Yes |
| | | 14.2 | Describe how non-independence has been handled | Yes |
| | | 14.3 | Justify decisions made | Yes |
| Meta-regression and model selection | 100% | 15.1 | Provide a rationale for the inclusion of moderators (covariates) that were evaluated in meta-regression models | Yes |
| | | 15.2 | Justify the number of parameters estimated in models, in relation to the number of effect sizes and studies (e.g. interaction terms were not included due to insufficient sample sizes) | Yes |
| | | 15.3 | Describe any process of model selection | NA |
| Publication bias and sensitivity analysis | 100% | 16.1 | Describe assessments of the risk of bias due to missing results (e.g. publication, time-lag, and taxonomic biases) | Yes |
| | | 16.2 | Describe any steps taken to investigate the effects of such biases (if present) | Yes |
| | | 16.3 | Describe any other analyses of robustness of the results, e.g. due to effect size choice, weighting or analytical model assumptions, inclusion or exclusion of subsets of the data, or the inclusion of alternative moderator variables in meta-regressions | Yes |

| | | | | |
|---|---|---|---|---|
| Clarification of post hoc analyses | 100% | 17.1 | When hypotheses were formulated after data analysis, this should be acknowledged. | Yes |
| Metadata, data, and code | | 18.1 | Share metadata (i.e. data descriptions) | Yes |
| | | 18.2 | Share data required to reproduce the results presented in the manuscript | Yes |
| | | 18.3 | Share additional data, including information that was not presented in the manuscript (e.g. raw data used to calculate effect sizes, descriptions of where data were located in papers) | Yes |
| | | 18.4 | Share analysis scripts (or, if a software package with graphical user interface (GUI) was used, then describe full model specification and fully specify choices) | Yes |
| Results of study selection process | 100% | 19.1 | Report the number of studies screened | Yes |
| | | 19.2 | Report the number of studies excluded at each stage of screening | Yes |
| | | 19.3 | Report brief reasons for exclusion from the full text stage | Yes |
| | | 19.4 | Present a Preferred Reporting Items for Systematic Reviews and Meta-Analyses (PRISMA)-like flowchart (www.prisma-statement.org). | Yes |
| Sample sizes and study characteristics | 75% | 20.1 | Report the number of studies and effect sizes for data included in meta-analyses | Yes |

| | | | | |
|---|---|---|---|---|
| | | 20.2 | Report the number of studies and effect sizes for subsets of data included in meta-regressions | Yes |
| | | 20.3 | Provide a summary of key characteristics for reported outcomes (either in text or figures; e.g. one quarter of effect sizes reported for vertebrates and the rest invertebrates) | Yes |
| | | 20.4 | Provide a summary of limitations of included moderators (e.g. collinearity and overlap between moderators) | NA |
| | | 20.5 | Provide a summary of characteristics related to individual study quality (risk of bias) | No |
| Meta-analysis | | 21.1 | Provide a quantitative synthesis of results across studies, including estimates for the mean effect size, with confidence/credible intervals | Yes |
| Heterogeneity | 100% | 22.1 | Report indicators of heterogeneity in the estimated effect (e.g. I2, tau2 and other variance components) | Yes |
| Meta-regression | | 23.1 | Provide estimates of meta-regression slopes (i.e. regression coefficients) and confidence/credible intervals | Yes |
| | | 23.2 | Include estimates and confidence/credible intervals for all moderator variables that were assessed (i.e. complete reporting) | Yes |
| | | 23.3 | Report interactions, if they were included | Yes |

| | | | | |
|---|---|---|---|---|
| | | 23.4 | Describe outcomes from model selection, if done (e.g. R2 and AIC) | NA |
| Outcomes of publication bias and sensitivity analysis | 100% | 24.1 | Provide results for the assessments of the risks of bias (e.g. Egger's regression, funnel plots) | Yes |
| | | 24.2 | Provide results for the robustness of the review's results (e.g. subgroup analyses, meta-regression of study quality, results from alternative methods of analysis, and temporal trends) | Yes |
| Discussion | 100% | 25.1 | Summarise the main findings in terms of the magnitude of effect | Yes |
| | | 25.2 | Summarise the main findings in terms of the precision of effects (e.g. size of confidence intervals, statistical significance) | Yes |
| | | 25.3 | Summarise the main findings in terms of their heterogeneity | Yes |
| | | 25.4 | Summarise the main findings in terms of their biological/practical relevance | Yes |
| | | 25.5 | Compare results with previous reviews on the topic, if available | Yes |
| | | 25.6 | Consider limitations and their influence on the generality of conclusions, such as gaps in the available evidence (e.g. taxonomic and geographical research biases) | Yes |
| Contributions and funding | 100% | 26.1 | Provide names, affiliations, and funding sources of all co-authors | Yes |

| | | | | |
|---|---|---|---|---|
| | | 26.2 | List the contributions of each co-author | Yes |
| | | 26.3 | Provide contact details for the corresponding author | Yes |
| | | 26.4 | Disclose any conflicts of interest | Yes |
| References | 100% | 27.1 | Provide a reference list of all studies included in the systematic review or meta-analysis | Yes |
| | | 27.2 | List included studies as referenced sources (e.g. rather than listing them in a table or supplement) | Yes |

**Supplementary Information S3:** The full keyword search string used, with a minor syntax edit at the beginning of the string for the electronic search engine Web of Science[†] (“TS =…”) and Scopus (“ALL…”), was:

(("maternal effect*" OR "parental effect*" OR "maternal invest*" OR "parental invest*" OR "maternal program*" OR "parental program*" OR "maternal hormon*" OR "epigenetic effect*" OR "epigenetic inheritance" OR "trans-generational phenotypic plasticity" OR "trans-generational plasticity" OR "transfer* from mother* to offspring" OR "transfer* from parent* to offspring" OR "transgeneration* induc*" OR "transgenerational adaptive plasticity" OR "transgenerational induction" OR "transgenerational phenotypic plasticity" OR "transgenerational plasticity" OR "transmit* between generations" OR "transmitted from mother* to offspring" OR "transmitted from parent* to offspring" OR "*generation* effect*" OR "anticipatory effect*" OR "bet hedging" OR "carry-over" OR "carryover" OR "developmental conditioning" OR "developmental programming" OR "fetal programming" OR "maternal conditioning" OR "maternal match*" OR "paternal conditioning" OR "silver spoon" OR "transgenerational process*" OR "*cross generation* effect" OR "imprinting" OR "inter-generational process*" OR "adaptive cross-generational hypothesis" OR "adaptive epigenetic memory" OR "between generation* effect*" OR "embryo* conditioning" OR "environmental matching hypothes*" OR "intergenerational phenotypic plasticity" OR "intergenerational plasticity" OR "maternal hormon* effect*" OR "maternal environment enhanc* offspring" OR "maternal environment* effect*" OR "maternal mediation hypothes*" OR "maternal mismatch hypothes*" OR "maternal rearing condition*" OR "mother-to-offspring transfer*" OR "mothers forewarn offspring" OR "parent-to-offspring transfer*" OR "parental environment* effect*" OR "parental rearing condition*" OR "paternal environment* effect*" OR "phenotypeenvironment *matching" OR "phenotype environment *match" OR "plasticity *cross generation*" OR "prim* the* offspring" OR "prime* offspring" OR "telescoping generation*" OR "priming" OR "generational memory") AND ("yolk steroid* hormon*" OR " yolk androgen*" OR "yolk androstenedion*" OR "yolk testosteron*" OR "yolk 5α-dihydrotestosteron*" OR "yolk DHT" OR "yolk glucocorticoid*" OR "yolk corticosteron*" OR "yolk cortisol" OR "yolk progesteron*" OR "yolk estrogen*" OR "yolk T4" OR "yolk T3" OR "yolk thyroid" OR "egg steroid* hormon*" OR "egg androgen*" OR "egg androstenedion*" OR "egg testosteron*" OR "egg 5α-dihydrotestosteron*" OR "egg DHT" OR "egg glucocorticoid*" OR "egg corticosteron*" OR "egg cortisol" OR "egg progesteron*" OR "egg estrogen*" OR "egg T4" OR "egg T3" OR "egg thyroid" OR "egg cort*" OR "yolk cort*" OR "egg estradiol" OR "yolk estradiol" OR "egg GC*" OR "yolk GC*" OR "egg hormon*" OR "yolk hormon*" OR "album* steroid* hormon*" OR "album* androgen*" OR "album* androstenedion*" OR " album* testosteron*" OR " album* 5α-dihydrotestosteron*" OR "album* DHT" OR "album* glucocorticoid*" OR "album* corticosteron*" OR "album* cortisol" OR "album* progesteron*" OR "album* estrogen*" OR "album* T4" OR "album* T3" OR "album* thyroid" OR "album* cort*" OR "album* estradiol" OR "album* GC*" OR "album* hormon*") AND ("fitness" OR "breed* success*" OR "clutch*" OR "hatch*" OR "fledg*" OR "embryo* development*" OR "reproductive success" OR "surviv*" OR "condition" OR "size" OR "growth" OR "mass" OR "tarsi" OR "tarsus" OR "wing*" OR "quality" OR "weight" OR "bone*" OR "standard length" OR "total length"))

[†] Databases covered with Web of Science Core Collection: Science Citation Index Expanded (SCIE); Social Sciences Citation Index (SSCI); Arts & Humanities Citation Index (AHCI); Emerging Sources Citation Index (ESCI)).

**Supplementary Information S4:** Full list of fitness proxies used in the current meta-analysis. Note that for some fitness proxies, it is difficult to fully disentangle maternal and offspring fitness (e.g., hatching or fledging success; discussed by Wolf & Wade 2001). In such cases, following the recommendations of Wolf & Wade (2001), we took a conservative approach, assigning to the mother those fitness proxies measured in the offspring that are largely or wholly controlled by her (e.g., hatching success, which is strongly influenced by maternal factors such as egg size; meta-analysis by Krist 2011). k: number of effect sizes; N: number of studies. Fitness proxies shown in grey were initially extracted but could not be included in the final analysis due to incomplete data.

¥ Lifetime number of sex-specific recruits produced by a female.

* Fitness proxies for offspring when recaptured as breeding adults.

| | Maternal fitness proxies | k | N | Offspring fitness proxies | k | N |
|---|---|---|---|---|---|---|
| 1 | Clutch size | 31 | 12 | Offspring survival | 13 | 8 |
| 2 | Egg mortality | 2 | 1 | Beak flank width | 10 | 1 |
| 3 | Hatching probability | 2 | 2 | Culmen | 5 | 1 |
| 4 | Hatching success | 42 | 33 | Fledgling number | 9 | 3 |
| 5 | Hatching failure | 1 | 1 | Fledging success | 14 | 10 |
| 6 | Maternal lifetime reproductive success | 2 | 1 | Fledging success (fledglings/hatchings) | 1 | 1 |
| 7 | Maternal longevity | 2 | 1 | Flipper length | 1 | 1 |
| 8 | Maternal recruitment¥ | 4 | 1 | Gape width | 15 | 2 |
| 9 | Maternal survival | - | - | Growth (mass) | 3 | 1 |
| 10 | Maternal number of reproductive events over different years | - | - | Growth (tarsus) | 3 | 1 |
| 11 | | | | Growth (PC1) | 2 | 1 |
| 12 | | | | Growth (PC2) | 2 | 1 |
| 13 | | | | Growth rate | 1 | 1 |
| 14 | | | | Growth rate (flipper length) | 1 | 1 |
| 15 | | | | Growth rate (mass) | 5 | 3 |
| 16 | | | | Growth rate (tarsus) | 1 | 1 |
| 17 | | | | Clutch size* | 2 | 1 |
| 18 | | | | Hatching number* | 11 | 6 |
| 19 | | | | Offspring survival years | 2 | |
| 20 | | | | Head-bill length | 1 | 1 |
| 21 | | | | Offspring mortality | 2 | 1 |
| 22 | | | | Pre-fledging survival probability | 4 | 1 |
| 23 | | | | Head length | 1 | 1 |

| | | | | | | |
|---|---|---|---|---|---|---|
| 24 | | | | Mass | 134 | 30 |
| 25 | | | | Mass gain | 2 | 1 |
| 26 | | | | Offspring recruitment | 1 | 1 |
| 27 | | | | Structural body size (mass and bill length) | 1 | 1 |
| 28 | | | | Structural body size (mass and tarsus) | 5 | 4 |
| 29 | | | | Tarsus length | 74 | 21 |
| 30 | | | | Wing length | 26 | 7 |
| 31 | | | | Fledging probability | - | - |
| 32 | | | | Fledgling success_egg1 | - | - |
| 33 | | | | Fledgling success_egg2 | - | - |
| 34 | | | | Fledgling success_egg3 | - | - |
| 35 | | | | Fledgling success_egg4 | - | - |
| 36 | | | | Fledgling success_egg5 | - | - |
| 37 | | | | Fledgling success_egg6 | - | - |
| 38 | | | | Growth rate (head length) | - | - |
| 39 | | | | Growth rate (wing) | - | - |
| 40 | | | | Survival | - | - |
| 41 | | | | Tail length | - | - |

**Supplementary Information S5:** Standardized email template sent to those authors whose articles had incomplete information to extract effect sizes and moderator information.

Dear XXX,

My name is Lucia Mentesana. I am a postdoctoral researcher at the Max Planck for Biological Intelligence (Germany). Together with Prof. Dr. Michaela Hau, Dr. Nicolas Adreani, Dr. Pietro B. D'Amelio & Dr. Alfredo Sánchez-Tójar, we are currently conducting a meta-analysis to test to what extent prenatal maternal egg hormone deposition relates to fitness in free-living birds. For this, we conducted a systematic literature search that identified 77 articles with relevant data for our meta-analysis and from where we have extracted all available information. Among these valuable studies, we found your work!

The reason why I am contacting you is because after carefully reading your work, we realized that we would need some extra information about your work to be able to include it in our meta-analysis (see details in the Word files attached). We were wondering if you could help us obtaining the missing information so that we can include your important work in our meta-analysis and make the results of it more robust, complete and generalizable. We would then of course make sure to cite your work in our manuscript.

We are aiming to finalize our analyses in about a month, so please, if you think you can provide us the missing information, but may need some extra time, do let us know. Please, do not hesitate to contact us if you have any questions.

Looking forward to hearing from you!
Thank you very much in advance!
Warm regards,
Lucia Mentesana et al.

**Supplementary Information S6:** Information on the continuous and categorical moderators used in the meta-regression models.

**Table S6.1:** Description of the continuous moderator used for meta-regression models.

| Continuous moderator | Description |
|---|---|
| Offspring relative age | Age of the offspring/Average nesting time across nests (both values obtained from information reported in the paper), with 0 = hatching day. median = 0.500; range = 0 - 1.125; mean = 0.448, SD = 0.302; k = 218; N = 19 |
| Square root of the inverse of the sample size | This is simply the square root of the inverse of the sample size used to calculate the effect size and its sampling variance. This moderator was used to test for small-study effects following (Nakagawa *et al.* 2022) |
| Year of publication (mean-centered) | The year when the study was published - the mean year of publication across our dataset. Mean-centering was performed to aid interpretation |

**Table S6.2:** Description of the categorical moderators used for meta-regression models.

| Categorical moderator | Moderator levels | Description |
|---|---|---|
| Study type | 1. Correlational<br>2. Experimental | Type of study |
| Egg hormone | 1. Androgens (androstenedione, 5α-dihydrotestosterone & testosterone)<br>2. Glucocorticoids<br>3. Thyroids (TH3 & TH4) | Type of hormone measured/manipulated |
| Experimental subject | 1. Mother<br>2. Egg | Subject that was experimentally manipulated |
| Hormone matrix | 1. Albumen<br>2. Yolk | Egg site where the hormone was measured |
| Experimental dose | 1. Supraphysiological<br>2. Within natural range | Range of hormone dose used for the experiment as stated by authors |
| Egg sampling method | 1. Entire egg removed<br>2. Biopsy | Method of matrix collection |
| Offspring life history stage | 1. Hatching<br>2. Before independence<br>3. After independence | Offspring age categories |
| Offspring sex | 1. Female<br>2. Male | Sex of the offspring determined in the paper |
| Data reporting | 1. Complete<br>2. Incomplete | Incomplete whenever authors did not report sample sizes, test for directionality, insufficient information on statistical models, insufficient information on statistical inference (e.g., missing p-values, CrI or CI) |
| Partial or selective reporting | 1. Yes<br>2. No | No: when authors did not report or partially reported results, for example, because the results were not statistically significant |

**Supplementary Information S7:** Effect size calculation.

**Table S7:** Equations used for converting the reported results into Pearson correlation coefficients ($r$; from Lajeunesse 2013; Nakagawa & Cuthill 2007). The sampling variance of $r$ was calculated as $Vr = (1 - r^2)^2 \div (N - 1)$ (Borenstein *et al.* 2009).

| Statistical test reported in the original paper | Equation |
|---|---|
| Absolute value | $r = [(Nexp\_survived * Ncontrol\_not\_survived) - (Ncontrol\_survived * Nexp\_not\_survived)] \div sqrt[(Nexp\_survived + Ncontrol\_survived) * (Nexp\_not\_survived + Ncontrol\_not\_survived) * (Nexp\_survived + Nexp\_not\_survived) * (Ncontrol\_survived + Ncontrol\_not\_survived)]$ |
| Chi-square value $(X^2)$ | $r = sqrt(X^2 \div N)$ |
| $F$-value | $r = sqrt(F \div (F + N – 2))$ |
| Spearman's rho correlation coefficient ($\rho$) | $r = 2 * sin(\pi\rho \div 6)$ |
| $t$-value | $r = t \div sqrt(t^2 + N – 2)$ |
| $z$-score | $r = z \div sqrt(N)$ |

**Supplementary Information S8:** Sensitivity analyses conducted to confirm the robustness of our results using a vector of sampling variances instead of the variance-covariance matrix. These analyses were performed for the intercept-only model of i) the entire database and ii) only offspring fitness traits, and iii) the hormone-type meta-regression for the offspring database. [*non-pre-registered analyses*]

| Models | Mean | L 95% CI | U 95% CI | L 95% PI | U 95% PI | Q | p-value | k | N |
|---|---|---|---|---|---|---|---|---|---|
| Intercept only models | | | | | | | | | |
| i) For the entire database | -0.051 | -0.259 | 0.157 | -0.766 | 0.663 | 24453 | 0.628 | 438 | 57 |
| ii) For the offspring database | -0.040 | -0.302 | 0.222 | -0.771 | 0.691 | 11713 | 0.763 | 352 | 42 |
| iii) Meta-regression model for the offspring database | | | | | | | | | |
| Androgens | -0.055 | -0.321 | 0.212 | -0.791 | 0.682 | | 0.688 | 260 | 33 |
| Glucocorticoids | 0.002 | -0.277 | 0.281 | -0.739 | 0.744 | 11662 | 0.986 | 56 | 8 |
| Thyroids | -0.018 | -0.303 | 0.267 | -0.761 | 0.726 | | 0.902 | 36 | 3 |

CI: confidence interval; PI: prediction interval; L: lower; U: upper; *Q:* heterogeneity; k: number of effect sizes; N: number of studies.

**Supplementary Information S9:** Sensitivity analyses testing the robustness of our results to different choices of effect size types. Note that these analyses used data on both maternal and offspring fitness.

**Table S9:** Model result table comparing the estimates from the original model (i.e., intercept-only model analyzing both Pearson and Biserial correlations) to those of four intercept-only models only including: (a) Pearson's *r* correlations, (b) Fisher's Zr, (c) Biserial correlations, or (d) Log-response ratio coefficients (lnRR). [*non-pre-registered analyses*]

| | Mean | L 95% CI | U 95% CI | L 95% PI | U 95% PI | $I^2$ total | p-value | k | N |
|---|---|---|---|---|---|---|---|---|---|
| Original (full) model | | | | | | | | | |
| | -0.072 | -0.330 | 0.186 | -0.851 | 0.707 | 95.3 | 0.233 | 438 | 57 |
| Sensitivity analysis: four intercept-only models | | | | | | | | | |
| a) Pearson *r* | 0.041 | -0.027 | 0.109 | -0.327 | 0.409 | 87.5 | 0.229 | 166 | 40 |
| b) Fisher's *Zr* | 0.008 | -0.030 | 0.046 | -0.227 | 0.243 | 73.2 | 0.670 | 166 | 40 |
| c) Biserial | -0.080 | -0.350 | 0.189 | -0.960 | 0.799 | 95.0 | 0.557 | 272 | 38 |
| d) lnRR | 0.004 | -0.004 | 0.011 | -0.070 | 0.077 | 98.4 | 0.359 | 269 | 37 |

CI: confidence interval; PI: prediction interval; L: lower; U: upper; $I^2$: total relative heterogeneity; k: number of effect sizes; N: number of studies.

Model a) "Pearson" used the subset of the data containing Pearson's *r* correlations that were either extracted directly from the original articles or transformed either from other types of correlations or from inferential statistics, and Model b) contains the "Fisher's Zr" transformations of those Pearson's r. Model c) "biserial" and d) "lnRR" used the subset of the data that were extracted as means, SDs and sample sizes, and then used for calculating either biserial correlations or lnRR (more in the section "2.6. Effect size calculation" from the main manuscript). lnRR cannot be calculated for non-ratio scale data, explaining the reduction in sample size in (d) compared to (c). None of the four sensitivity analyses deviated from the conclusion of the main model, that is, the meta-analytic mean is non-statistically significantly different from zero. However, note that these models do not differentiate between hormone groups, and that a) the full model contains both experimental and correlational data, whereas the subset of data used for c) and d) corresponds to results from experimental data only.

**Supplementary Information S10:** Sensitivity analyses testing the robustness of our results for predictions BP.1.1 - BP.1.3. Note that these analyses used data on offspring fitness only.

**Table S10.1:** Model result table showing the estimates from three intercept-only models including the data of each egg hormone separately compared to the estimates shown in the main text, which come from a meta-regression including hormone type as a moderator with three levels.

| Models | Mean | L 95% CI | U 95% CI | L 95% PI | U 95% PI | *Q* | p-value | k | N |
|---|---|---|---|---|---|---|---|---|---|
| BH.1.1 (androgens) | -0.066 | -0.439 | 0.307 | -1.008 | 0.876 | 21449 | 0.728 | 260 | 33 |
| BH.1.2 (glucocorticoids) | 0.077 | -0.032 | 0.186 | -0.225 | 0.378 | 176 | 0.164 | 56 | 8 |
| BH.1.3 (thyroids) | 0.022 | -0.083 | 0.127 | -0.331 | 0.375 | 200 | 0.672 | 36 | 3 |

CI: confidence interval; PI: prediction interval; L: lower; U: upper; *Q:* heterogeneity; k: number of effect sizes; N: number of studies.

**Table S10.2:** Heterogeneity metrics and stratification from three intercept-only models including the data of each egg hormone separately.

| | Androgens | | | | Glucocorticoids | | | | Thyroids | | | |
|---|---|---|---|---|---|---|---|---|---|---|---|---|
| | $I^2$ *(%)* | *CVH2* | *M2* | $\sigma^2$ | $I^2$ *(%)* | *CVH2* | *M2* | $\sigma^2$ | $I^2$ *(%)* | *CVH2* | *M2* | $\sigma^2$ |
| Total | 95.88 | 44.20 | 0.98 | 0.19 | 74.52 | 3.34 | 0.77 | 0.02 | 80.84 | 57.08 | 0.98 | 0.03 |
| Among-study (Study ID) | 0.00 | 0.00 | 0.00 | 0.00 | 0.00 | 0.00 | 0.00 | 0.00 | 0.00 | 0.00 | 0.00 | 0.00 |
| Among-laboratory (Laboratory ID) | 0.00 | 0.00 | 0.00 | 0.00 | 42.17 | 1.89 | 0.44 | 0.01 | 0.00 | 0.00 | 0.00 | 0.00 |
| Among-population (Population ID) | 0.00 | 0.00 | 0.00 | 0.00 | 0.00 | 0.00 | 0.00 | 0.00 | 0.00 | 0.00 | 0.00 | 0.00 |
| Among-species (Species name) | 0.00 | 0.00 | 0.00 | 0.00 | 0.00 | 0.00 | 0.00 | 0.00 | 0.00 | 0.00 | 0.00 | 0.00 |
| Phylogenetic relationships (Species phylogeny) | 54.27 | 25.02 | 0.55 | 0.11 | 0.00 | 0.00 | 0.00 | 0.00 | 0.00 | 0.00 | 0.00 | 0.00 |
| Within-Study (Effect ID) | 41.61 | 19.18 | 0.42 | 0.08 | 32.35 | 1.45 | 0.33 | 0.01 | 80.84 | 57.08 | 0.98 | 0.03 |

$I^2$: variance-standardized (AKA relative heterogeneity), *CVH2*: mean-standardized heterogeneity, *M2*: variance-mean-standardized heterogeneity, $\sigma^2$: total heterogeneity.

**Table S10.3**. Small-study effects analyses for each of the three intercept-only models including the data of each egg hormone separately.

| Hormone | Intercept | L 95% CI | U 95% CI | p-value | Sqrt (1/N) | L 95% CI | U 95% CI | p-value | $R^2_{marginal}$ | k | N |
|---|---|---|---|---|---|---|---|---|---|---|---|
| Androgen | -0.136 | -0.451 | 0.179 | 0.397 | 0.375 | -0.441 | 1.192 | 0.367 | 0.5% | 333 | 46 |
| Glucocorticoids | 0.057 | -0.085 | 0.198 | 0.426 | 0.242 | -0.855 | 1.338 | 0.661 | 0.9% | 66 | 10 |
| Thyroids | -0.227 | -0.467 | 0.013 | 0.063 | 3.016 | 0.422 | 5.611 | 0.024 | 23.8% | 39 | 3 |

CI: confidence interval; PI: prediction interval; L: lower; U: upper; k: number of effect sizes; N: number of studies.

**Supplementary Information S11.** Effect that maternal egg hormones have on offspring fitness differs depending on the matrix in which the hormone was measured/manipulated (BEH.2).

To test if the effect of maternal egg hormones on offspring fitness differs depending on the matrix in which the hormone was measured/manipulated (BEH.2), we planned to fit a phylogenetic multilevel meta-regression with the same random effects structure as used in the other models (i.e, study ID, laboratory ID, population ID, species name, phylogeny, and effect ID) and 'hormone type' (levels: androgens, glucocorticoids, and thyroids), 'egg location' (levels: albumen, yolk), and their interaction as moderators. However, data was available only for glucocorticoids and three species (*Parus major*, *Larus michahellis*, and *Troglodytes aedon*). We therefore had to reduce our model to a multilevel meta-regression including as only three random effects (i.e., study ID, species name, and effect ID) and 'egg location' as the only. Given the small data available (k = 18, N = 4), the results of this meta-regression should be interpreted with extreme caution, which is the main reason for why these results are presented in the supplementary material despite BEH.2. being part of our original list of pre-registered exploratory hypothesis.

We found a statistically nonsignificant positive association between hormone concentrations and offspring fitness regardless of the egg site where the hormone was measured (albumen: r = 0.028, [95% CI = -0.164, 0.219], [95% PI = -0.270, 0.325], p-value = 0.772; yolk: r = 0.094, [95% CI = -0.100, 0.287], [95% PI = -0.205, 0.393], p-value = 0.332; $R^2$marginal = 8.1%; p-value albumen vs yolk = 0.625).

**Supplementary Information S12:** Publication bias tests and results.

**Publication bias analyses**

We assessed publication bias, both small-study and decline effects, following Nakagawa *et al.* (2022). All models employed the same random effect structure as reported for the analyses above. We used the complete database containing information on both maternal and offspring fitness. For small-study effects, we conducted a multilevel meta-regression, including the ‘square root of the inverse of the sample size’ of each effect size as a moderator. Additionally, we investigated temporal patterns in the data that could highlight the existence of decline effects (i.e., effect sizes decreasing over time; Sánchez-Tójar *et al.* 2018; Trikalinos & Ioannidis 2005) by conducting a multilevel meta-regression where the ‘year of publication’ was mean-centered and included as a moderator. Note that these two analyses using the entire dataset were not pre-registered, but we added them to provide an informative overall overview of small-study and decline effects. We also tested for the existence of small-study and decline effects for each hormone separately by rerunning the same two models described above but adding the interaction between the ‘square root of the inverse of the sample size’ or ‘year of publication’ and 'hormone type' (levels: androgens, glucocorticoids, and thyroids), respectively.

In addition, to test whether effect sizes are larger in studies that selectively or incompletely report statistically significant results, we conducted two meta-regression models. To test the effect of partial or selective reporting, we included 'partial or selective reporting' (levels: yes and no), 'hormone type,' and their interaction as moderators. To examine the effect of data reporting completeness, we initially planned to include 'data reporting completeness' (levels: complete and incomplete), 'hormone type,' and their interaction as moderators. However, due to insufficient data for glucocorticoids and thyroids, we could only test this for androgens (k = 333, N = 46), so our final model included 'data reporting completeness' as the only moderator. We also had pre-registered a meta-regression to test if blind data collection leads to smaller effect sizes as shown in previous studies (Freeberg *et al.* 2024; Holman *et al.* 2015; Keaney *et al.* 2024), but we could not perform this test because only 3 studies reported to have used a blind data collection protocol.

Lastly, we ran a multi-moderator multilevel meta-regression to estimate adjusted overall effects, and the percentage of heterogeneity explained by those moderators combined ($R^2_{marginal}$). For this, we mean-centered and included 'study type' as a moderator, as well as the interaction between the 'square root of the inverse of the sample size' and 'hormone type', and the interaction between 'year' and 'hormone type'.

**Publication bias results**

The evidence for the existence of overall small-study effects across the full dataset was statistically nonsignificant and the sign was contrary to our expectations (Figure S10A; intercept = -0.135, [95% CI = -0.436, 0.166], p-value = 0.380; slope = 0.414, [95% CI = -0.271,1.100], p-value = 0.236; k = 438, N = 57; $R^2_{marginal}$ = 0.6%), suggesting that positive effect sizes are associated with effect sizes of lower precision across the full dataset combining all three hormones. In other words, some negative effect sizes of lower precision likely remain unpublished in the scientific literature, and although adjusting for their absence would not seemingly change the statistical significance of the mean effect, it could result in a ca. 100% increase in its magnitude from -0.072 to -0.135. Our model testing for hormone-specific small-study effects did not show statistically significant evidence of small-study effects for any of the three hormones (Figures S10B-D; $R^2_{marginal}$ = 1.0%), but the tendency described above suggesting that some negative effect sizes of lower precision may remain unpublished was present for all three hormones (adjusted effects: androgens: *r* = -0.091, [95% CI = -0.374, 0.193], [95% PI = -0.906, 0.725], k = 333, N = 46; glucocorticoids: *r* = -0.030, [95% CI = -0.346, 0.286], [95% PI = -0.857, 0.797], k = 66, N = 10; thyroids: *r* = 0.033, [95% CI = -0.347, 0.414], [95% PI = -0.820, 0.887], k = 39, N = 3; Table S10.1).

We also found no clear evidence of decline effects either when combining all hormones together (intercept = -0.068, [95% CI = -0.326, 0.190]; slope = 0.002, [95% CI = -0.007, 0.011]; k = 438, N = 57; $R^2_{marginal}$ = 0.1%), or when testing for hormone-specific patterns via meta-regression ($R^2_{marginal}$ = 0.4%; Table S10.2). The adjusted hormone-specific effects from the latter model remained statistically nonsignificant (androgens: *r* = -0.082, [95% CI = -0.346, 0.183], [95% PI = -0.868, 0.705], k = 333, N = 46; glucocorticoids: *r* = -0.039, [95% CI = -0.322, 0.245], [95% PI = -0.832, 0.754], k = 66, N = 10 ; thyroids: *r* = -0.012, [95% CI = -0.612, 0.588], [95% PI = -0.965, 0.941], k = 39, N = 3).

We were able to categorize whether researchers did or did not selectively report data for only 38.1% of effect sizes (k = 168, N = 25). Authors partially or selectively reported data in 83 cases, while in 85 cases they did not. For thyroid hormones, but not for androgens or glucocorticoids, we found evidence that selectively and/or partially reported results influenced the effect size (Table S10.3). However, these results should be interpreted with extreme caution, as only one study (k = 10) on thyroid hormones selectively/partially reported outcomes, making it impossible to determine whether the observed effect size is due to reporting biases or the study's nature. The moderator explained a relatively high amount of heterogeneity ($R^2_{marginal}$ = 18.1%) and, within each hormone group, the two levels (i.e., no *vs* yes partial/selective reporting) did not differ statistically from each other (androgens: p-value = 0.624; glucocorticoids: p-value = 0.121; thyroids: p-value = 0.369).

We were able to examine the effect of data reporting completeness only for androgens. We found no evidence of an effect of data reporting completeness (complete: $r$ = -0.081, [95% CI = -0.344, 0.182], [95% PI = -0.912, 0.751], k = 299, N = 45; incomplete: $r$ = -0.040, [95% CI = -0.331, 0.251], [95% PI = -0.881, 0.801], k = 34, N = 8; p-value complete *vs* incomplete = 0.561) and this moderator explained little variance ($R^2_{marginal}$ = 0.1%). However, our results should be interpreted cautiously because they do not account for the fact that 221 potentially suitable effect sizes (33.5% of 659) could not be included in our final dataset due to incomplete reporting information.

Last, the multi-moderator model that we originally planned to estimate the total amount of heterogeneity explained by complete-case moderators, only contained a test for both small-study and decline effects simultaneously due to the lack of complete-case moderators in our dataset. This multi-moderator model resulted in similar results as those of the uni-moderator model testing for publication bias ($R^2_{marginal}$ = 1.0%; Table S10.4).

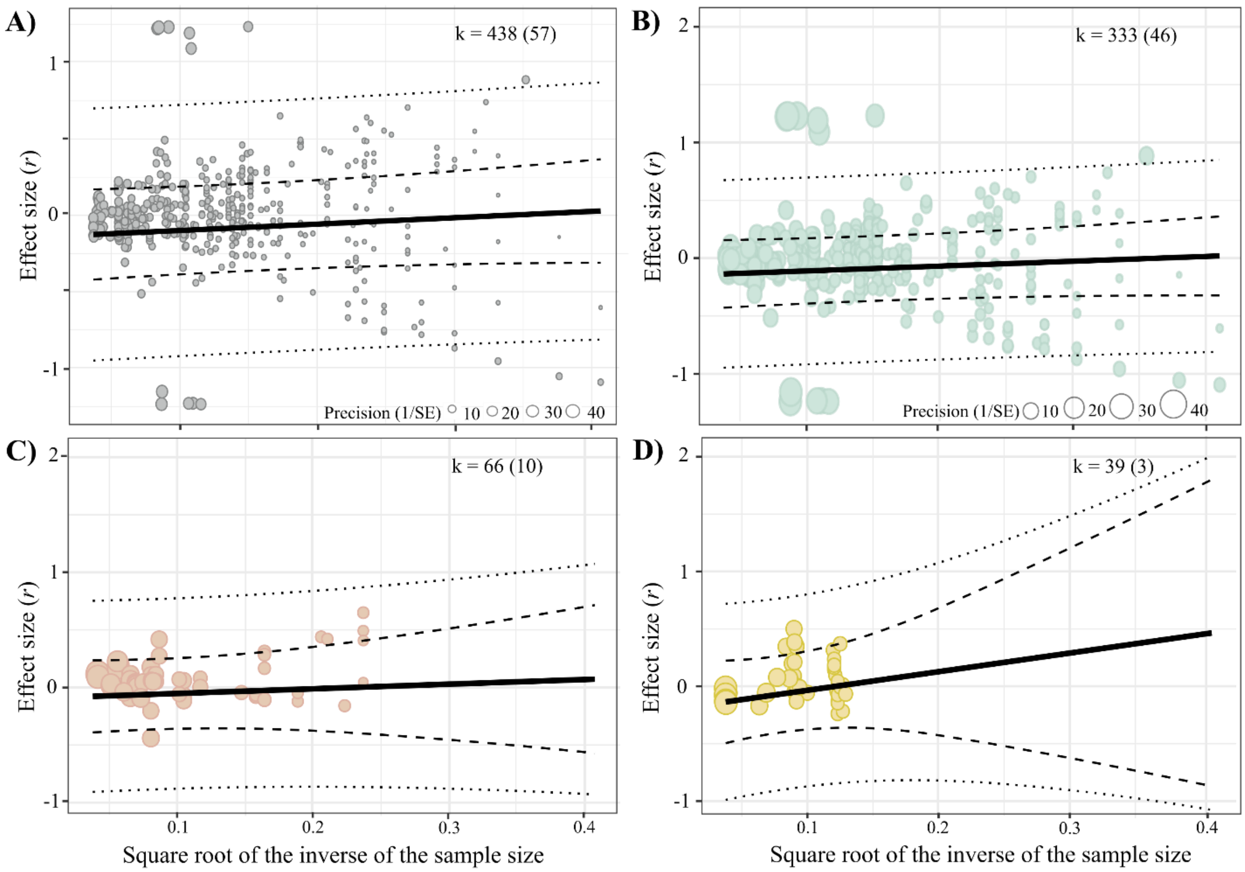


**Figure S12:** Orchard plots showing small-study effects biases in our dataset. A) For all studies, larger positive effect sizes were associated with effect sizes of lower prediction. For B) androgens, C) glucocorticoids, and D) thyroids no clear association was present. k is the number of individual effect sizes, the number of studies is shown in brackets.

**Table S12.1:** Hormone-specific small-study effects analyses across the full dataset (i.e., considering both maternal and offspring fitness).

| | Mean | SE | t-value | df | p-value | L 95% CI | U 95% CI |
|---|---|---|---|---|---|---|---|
| Intercept (Androgens) | -0.150 | 0.155 | -0.966 | 432 | 0.334 | -0.455 | 0.155 |
| Glucocorticoids | 0.066 | 0.104 | 0.633 | 432 | 0.527 | -0.139 | 0.270 |
| Thyroids | -0.058 | 0.200 | -0.288 | 432 | 0.774 | -0.451 | 0.336 |
| sqrt(1/N) | 0.427 | 0.384 | 1.110 | 432 | 0.267 | -0.329 | 1.182 |
| Glucocorticoids * sqrt(1/N) | -0.041 | 0.953 | -0.043 | 432 | 0.966 | -1.915 | 1.833 |
| Thyroids * sqrt(1/N) | 1.306 | 2.091 | 0.624 | 432 | 0.533 | -2.805 | 5.417 |

df: degrees of freedom; L: lower; U: upper.

**Table S12.2:** Decline effects analyses across the full dataset (i.e., considering both maternal and offspring fitness).

| | Mean | SE | t-value | df | p-value | L 95% CI | U 95% CI |
|---|---|---|---|---|---|---|---|
| Intercept (Androgens) | -0.082 | 0.135 | -0.606 | 432 | 0.545 | -0.346 | 0.183 |
| Glucocorticoids | 0.043 | 0.066 | 0.656 | 432 | 0.512 | -0.086 | 0.172 |
| Thyroids | 0.070 | 0.274 | 0.253 | 432 | 0.800 | -0.470 | 0.609 |
| Year | -0.000 | 0.005 | -0.004 | 432 | 0.997 | -0.010 | 0.010 |
| Glucocorticoids * Year | 0.005 | 0.009 | 0.513 | 432 | 0.608 | -0.014 | 0.023 |
| Thyroids * Year | -0.003 | 0.050 | -0.068 | 432 | 0.946 | -0.101 | 0.094 |

df: degrees of freedom; L: lower; U: upper.

**Table S12.3:** Results of the phylogenetic multilevel meta-regression models testing whether the associations between hormone concentrations and both maternal and offspring fitness differ between effect sizes from studies categorized as partially or selectively reporting results.

| | Partial or selective reporting | Mean | L 95% CI | U 95% CI | L 95% PI | U 95% PI | p-value | k | N |
|---|---|---|---|---|---|---|---|---|---|
| Androgens | No | -0.020 | -0.083 | 0.042 | -0.303 | 0.263 | 0.528 | 56 | 7 |
| | Yes | -0.004 | -0.070 | 0.077 | -0.282 | 0.290 | 0.919 | 54 | 12 |
| Glucocorticoids | No | 0.091 | -0.003 | 0.185 | -0.200 | 0.383 | 0.057 | 14 | 3 |
| | Yes | -0.022 | -0.133 | 0.090 | -0.320 | 0.276 | 0.704 | 19 | 5 |
| Thyroids | No | 0.095 | -0.133 | 0.324 | -0.263 | 0.454 | 0.410 | 15 | 1 |
| | Yes | 0.227 | 0.050 | 0.404 | -0.101 | 0.555 | 0.012 | 10 | 1 |

CI: confidence interval; PI: prediction interval; L: lower; U: upper; k = number of effect sizes; N = number of studies.

**Table S12.4:** Pseudo-all-in model to estimate the total amount of heterogeneity explained by complete-case moderators (i.e., small-study and decline effects). Note that these analyses used data on both maternal and offspring fitness.

| | Mean | SE | t-value | df | p-value | L 95% CI | U 95% CI |
|---|---|---|---|---|---|---|---|
| Intercept (Androgens) | -0.148 | 0.158 | -0.934 | 429 | 0.351 | -0.459 | 0.163 |
| Glucocorticoids | 0.053 | 0.109 | 0.484 | 429 | 0.628 | -0.161 | 0.266 |
| Thyroids | -0.220 | 0.435 | -0.506 | 429 | 0.613 | -1.074 | 0.634 |
| sqrt(1/N) | 0.444 | 0.394 | 1.126 | 429 | 0.261 | -0.331 | 1.219 |
| Year | 0.001 | 0.005 | 0.177 | 429 | 0.859 | -0.010 | 0.012 |
| Glucocorticoids * sqrt(1/N) | -0.018 | 0.964 | -0.019 | 429 | 0.985 | -1.912 | 1.877 |
| Thyroids * sqrt(1/N) | 1.774 | 2.335 | 0.760 | 429 | 0.448 | -2.825 | 6.362 |
| Glucocorticoids * Year | 0.004 | 0.009 | 0.431 | 429 | 0.667 | -0.015 | 0.023 |
| Thyroids * Year | 0.022 | 0.056 | 0.397 | 429 | 0.692 | -0.087 | 0.132 |

df: degrees of freedom; L: lower; U: upper.

**Supplementary Information S13:** Heterogeneity metrics and stratification for a meta-analysis testing the association between maternal egg hormones and fitness - both offspring and maternal - across 19 bird species.

| | $I^2$ (%) | $CVH2$ | $M2$ | $\sigma^2$ |
|---|---|---|---|---|
| Total | 95.31 | 26.72 | 0.96 | 0.14 |
| Among-study (Study ID) | 0.00 | 0.00 | 0.00 | 0.00 |
| Among-laboratory (Laboratory ID) | 0.00 | 0.00 | 0.00 | 0.00 |
| Among-population (Population ID) | 0.00 | 0.00 | 0.00 | 0.00 |
| Among-species (Species name) | 0.00 | 0.00 | 0.00 | 0.00 |
| Phylogenetic relationships (Species phylogeny) | 37.67 | 10.56 | 0.38 | 0.06 |
| Within-Study (Effect ID) | 57.64 | 16.16 | 0.58 | 0.09 |

$I^2$: variance-standardized (AKA relative heterogeneity), $CVH2$: mean-standardized heterogeneity, $M2$: variance-mean-standardized heterogeneity, $\sigma^2$: total heterogeneity.

$I^2_{total}$ indicates that heterogeneity in our dataset is, on average, ca. 20 times larger than statistical noise, and $CVH2_{total}$ and $M2_{total}$ indicate that heterogeneity is, on average, ca. 27 times larger than the meta-analytic mean. These analyses also suggest that once the phylogenetic relationships between the 19 bird species included in our meta-analysis have been accounted for, the only important source of heterogeneity is the within-study level, indicating that it is within-study level predictors that are most likely driving the observed context-dependency for the hypothesis that maternal egg hormones associate with fitness. Said differently, despite the high levels of heterogeneity observed, the meta-analytic mean observed could still be generalized at the between-study level. if we define replication as the testing of the null hypothesis at the between-study level and when within-study variations (methodological and biological) can be accounted for.

**Supplementary Information S14:** Results of a phylogenetic multilevel meta-regression (without intercept) model testing the association of egg hormone concentrations on offspring fitness depending on hormone type (androgens, glucocorticoids, and thyroids)*. A graphical visualization of these results is presented in the main manuscript as Figure 3.

| | Mean | L 95% CI | U 95% CI | L 95% PI | U 95% PI | p-value | k | N |
|---|---|---|---|---|---|---|---|---|
| Androgens | -0.076 | -0.429 | 0.277 | -0.951 | 0.799 | 0.672 | 260 | 33 |
| Glucocorticoids | -0.023 | -0.388 | 0.343 | -0.903 | 0.857 | 0.903 | 56 | 8 |
| Thyroids | -0.026 | -0.408 | 0.356 | -0.913 | 0.861 | 0.892 | 36 | 3 |

* CI: confidence interval; PI: prediction interval; L: lower; U: upper; k = number of effect sizes; N = number of studies.

**Supplementary Information S15:** Estimates of three phylogenetic multilevel meta-regressions used to investigate the relationship between egg hormones and fitness in: A) correlational *vs* experimental studies, B) experiments that manipulated the mother's hormone concentrations *vs* the egg's hormone concentrations, and C) experiments that manipulated maternal egg hormones within *vs* above the natural physiological range§. A graphical visualization of these results is presented in the main manuscript as Figure 6.

| Fitness effects | | | | | | | | | |
|---|---|---|---|---|---|---|---|---|---|
| | | Mean | L 95% CI | U 95% CI | L 95% PI | U 95% PI | p-value | k | N |
| **A)** *Correlational vs experimental studies* | | | | | | | | | |
| Androgens | Correlational | -0.004 | -0.276 | 0.269 | -0.796 | 0.789 | 0.978 | 90 | 9 |
| | Experimental | -0.095 | -0.351 | 0.160 | -0.882 | 0.691 | 0.463 | 243 | 38 |
| Glucocorticoids | Correlational | 0.067 | -0.252 | 0.387 | -0.743 | 0.878 | 0.679 | 13 | 2 |
| | Experimental | -0.055 | -0.343 | 0.232 | -0.853 | 0.742 | 0.705 | 53 | 8 |
| **B)** *Experimental subject* | | | | | | | | | |
| Androgens | Mother | -0.136 | -0.591 | 0.297 | -1.074 | 0.780 | 0.535 | 15 | 3 |
| | Egg | -0.069 | -0.414 | 0.277 | -0.970 | 0.833 | 0.695 | 188 | 26 |
| Glucocorticoids | Mother | -0.055 | -0.491 | 0.382 | -0.994 | 0.885 | 0.806 | 17 | 3 |
| | Egg | -0.058 | -0.444 | 0.329 | -0.976 | 0.860 | 0.769 | 27 | 3 |
| **C)** *Experimental dose* | | | | | | | | | |
| Androgens | Natural range | -0.113 | -0.450 | 0.223 | -1.112 | 0.885 | 0.505 | 82 | 17 |
| | Supraphysiol. range | 0.188 | -0.343 | 0.718 | -0.891 | 1.267 | 0.484 | 7 | 2 |

§ CI: confidence interval; PI: prediction interval; L: lower; U: upper; k = number of effect sizes; N = number of studies.

**Supplementary Information S16:** Results of the phylogenetic multilevel meta-regression model for our entire dataset, with 'fitness proxies' as the only moderator. Following our pre-registration, from all possible fitness proxies (see Supplementary Information S2), we only included those levels that had at least 5 data points (15 levels, 19 bird species). Also, we included the same random effects as in the other meta-regression models: study ID, laboratory ID, population ID, species name, species phylogeny, and effect ID.

| Category | Fitness proxy reported in paper | Mean | L 95% CI | U 95% CI | L 95% PI | U 95% PI | p-value | k | N |
|---|---|---|---|---|---|---|---|---|---|
| Clutch size | | -0.033 | -0.319 | 0.254 | -0.814 | 0.749 | 0.822 | 33 | 13 |
| Hatching number | Hatching number | -0.001 | -0.309 | 0.308 | -0.791 | 0.789 | 0.996 | 13 | 7 |
| | Egg mortality | | | | | | | | |
| Hatching success | Hatching success | -0.167 | -0.430 | 0.095 | -0.940 | 0.606 | 0.212 | 45 | 35 |
| | Hatching probability | | | | | | | | |
| | Hatching failure | | | | | | | | |
| Beak flank width | | 0.027 | -0.309 | 0.362 | -0.774 | 0.828 | 0.876 | 10 | 1 |
| Culmen | | 0.298 | -0.151 | 0.746 | -0.557 | 1.152 | 0.192 | 5 | 1 |
| Gape width | | 0.016 | -0.291 | 0.323 | -0.773 | 0.805 | 0.918 | 15 | 2 |
| Mass | | -0.025 | -0.279 | 0.229 | -0.796 | 0.745 | 0.846 | 134 | 30 |
| Tarsus | | -0.040 | -0.298 | 0.218 | -0.812 | 0.732 | 0.761 | 74 | 21 |
| Wing | Wing | -0.053 | -0.331 | 0.224 | -0.832 | 0.725 | 0.706 | 27 | 8 |
| | Flipper length | | | | | | | | |
| Structural body size | Structural body size (mass and tarsus) | -0.031 | -0.386 | 0.323 | -0.840 | 0.778 | 0.863 | 6 | 4 |
| | Structural body size (mass and | | | | | | | | |

| | | | | | | | | | |
|---|---|---|---|---|---|---|---|---|---|
| | bill length) | | | | | | | | |
| Growth | Growth (PC1) | 0.052 | -0.238 | 0.342 | -0.731 | 0.835 | 0.726 | 20 | 9 |
| | Growth (PC2) | | | | | | | | |
| | Growth (mass) | | | | | | | | |
| | Growth (tarsus) | | | | | | | | |
| | Growth rate | | | | | | | | |
| | Growth rate (mass) | | | | | | | | |
| | Growth rate (tarsus) | | | | | | | | |
| | Growth rate (flipper length) | | | | | | | | |
| | Mass gain | | | | | | | | |
| Fledgling number | | 0.159 | -0.166 | 0.484 | -0.638 | 0.955 | 0.337 | 9 | 3 |
| Fledgling success | Fledgling success | -0.056 | -0.343 | 0.230 | -0.838 | 0.725 | 0.700 | 19 | 11 |
| | Pre-fledgling survival probability | | | | | | | | |
| | Fledgling success (fledglings/hatchings) | | | | | | | | |
| Offspring survival | Offspring survival | 0.032 | -0.263 | 0.327 | -0.753 | 0.817 | 0.931 | 15 | 9 |
| | Offspring mortality | | | | | | | | |
| Recruit | Offspring recruitment | -0.152 | -0.492 | 0.188 | -0.954 | 0.651 | 0.381 | 7 | 3 |
| | Female recruitment | | | | | | | | |
| | Offspring survival years | | | | | | | | |

CI: confidence interval; PI: prediction interval; L: lower; U: upper; k = number of effect sizes; N = number of studies.

**Supplementary Information S17:** Results of the phylogenetic multilevel meta-regression model for our entire dataset, with 'developmental model' (levels: altricial and precocial) as the only moderator. We included the same random effects as in the other meta-regression models: study ID, laboratory ID, population ID, species name, species phylogeny, and effect ID.

| | Mean | L 95% CI | U 95% CI | L 95% PI | U 95% PI | p-value | k | N |
|---|---|---|---|---|---|---|---|---|
| Altricial | -0.090 | -0.370 | 0.191 | -0.888 | 0.708 | 0.530 | 401 | 46 |
| Precocial | -0.018 | -0.399 | 0.362 | 0.857 | 0.820 | 0.925 | 37 | 11 |

CI: confidence interval; PI: prediction interval; L: lower; U: upper; k = number of effect sizes; N = number of studies.

### Supplementary Figures

**Supplementary Figure 1.** Quality assessment of our systematic review and meta-analysis testing the extent to which prenatal maternal hormone deposition into eggs relates to fitness in wild birds. For it, we filled out the ‘Interactive PRISMA-EcoEvo Checklist’ (https://prisma-ecoevo.shinyapps.io/checklist; O’Dea *et al.* 2021).

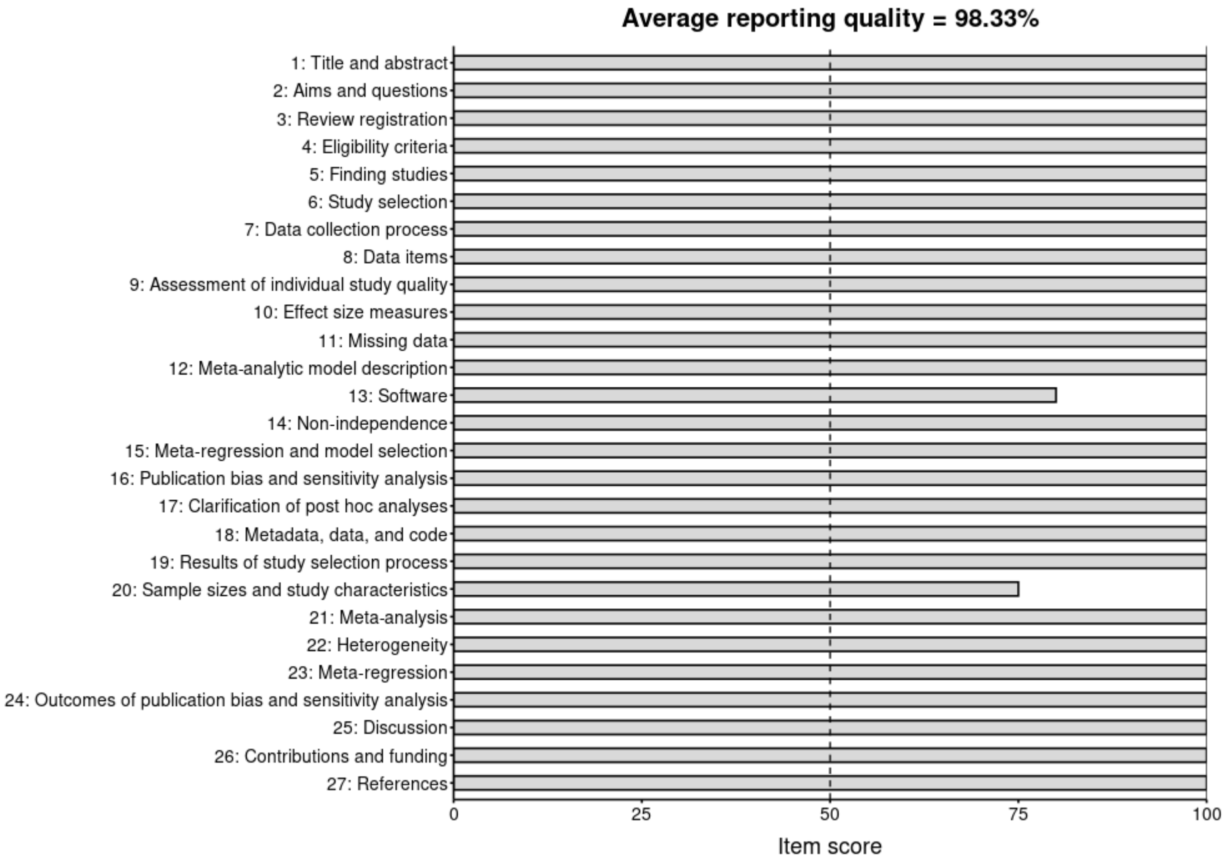

**Supplementary Figure 2.** Complete decision tree used for both title-and-abstract and full-text screening. Note that in our pre-registration (Mentesana *et al.* 2021), we had separate trees for title-and-abstract and full-text screening, but here we present a unified and complete version for clarity.

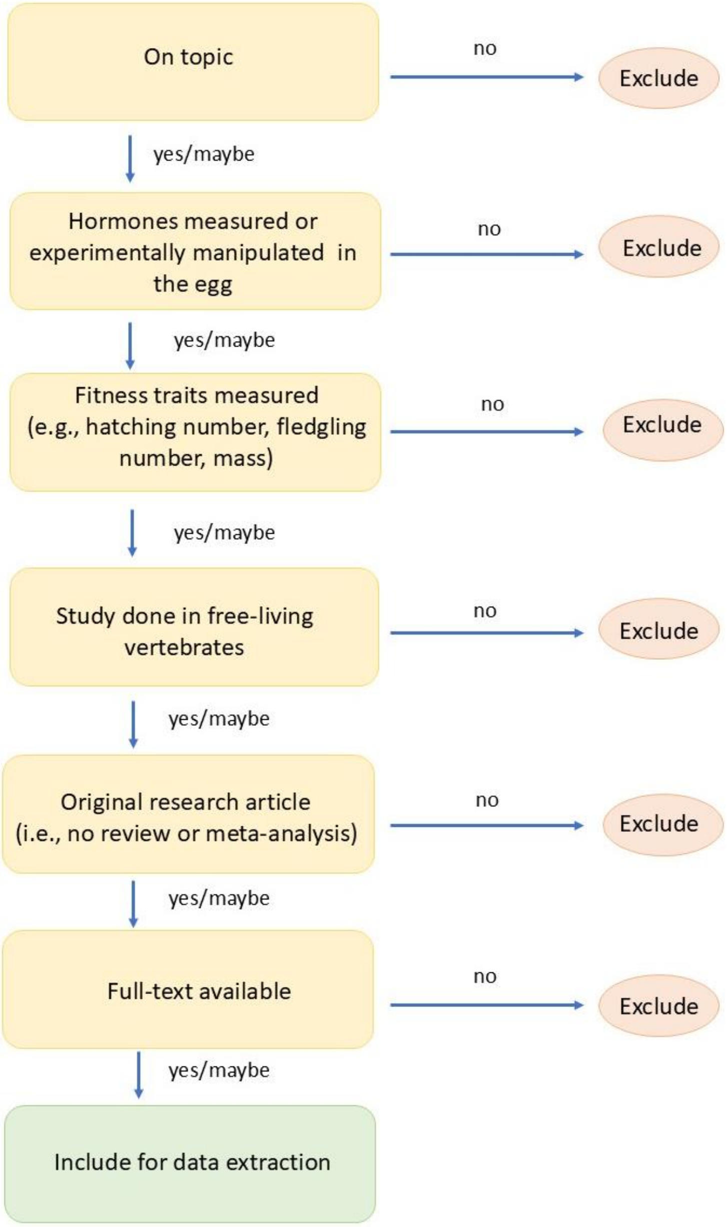

**Supplementary Figure 3.** PRISMA flow diagram summarizing our literature search. Of 80 eligible articles, we could not include 20 studies in the meta-analysis because articles reported incomplete information (e.g., missing sample sizes) and the authors did not provide such information despite our author contacting efforts (see “Materials and Methods” section), 1 article did not explore the effect of a hormone *per se* but that of a hormone blocker, and 2 studies were conducted in turtles.

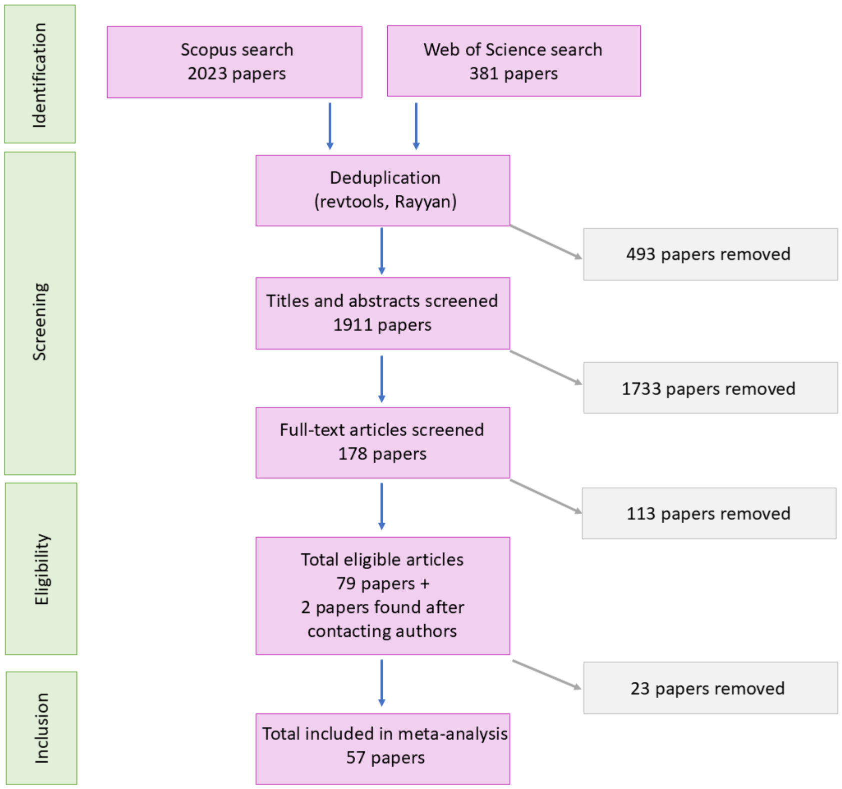

**Supplementary Figure 4.** Heterogeneity metrics and stratification for a meta-analysis testing the association between maternal egg hormones and fitness across 19 bird species. The heterogeneity is quantified using A) raw variance ($\sigma^2$), B) source measure $I^2$, C) magnitude measure $CV$, and D) magnitude measure $M$, and stratified at phylogenetic (Phylo), non-phylogenetic among-species (Spp), among-population (Pop), among-laboratory (Lab), among- (Among) and within-study (Within) levels.

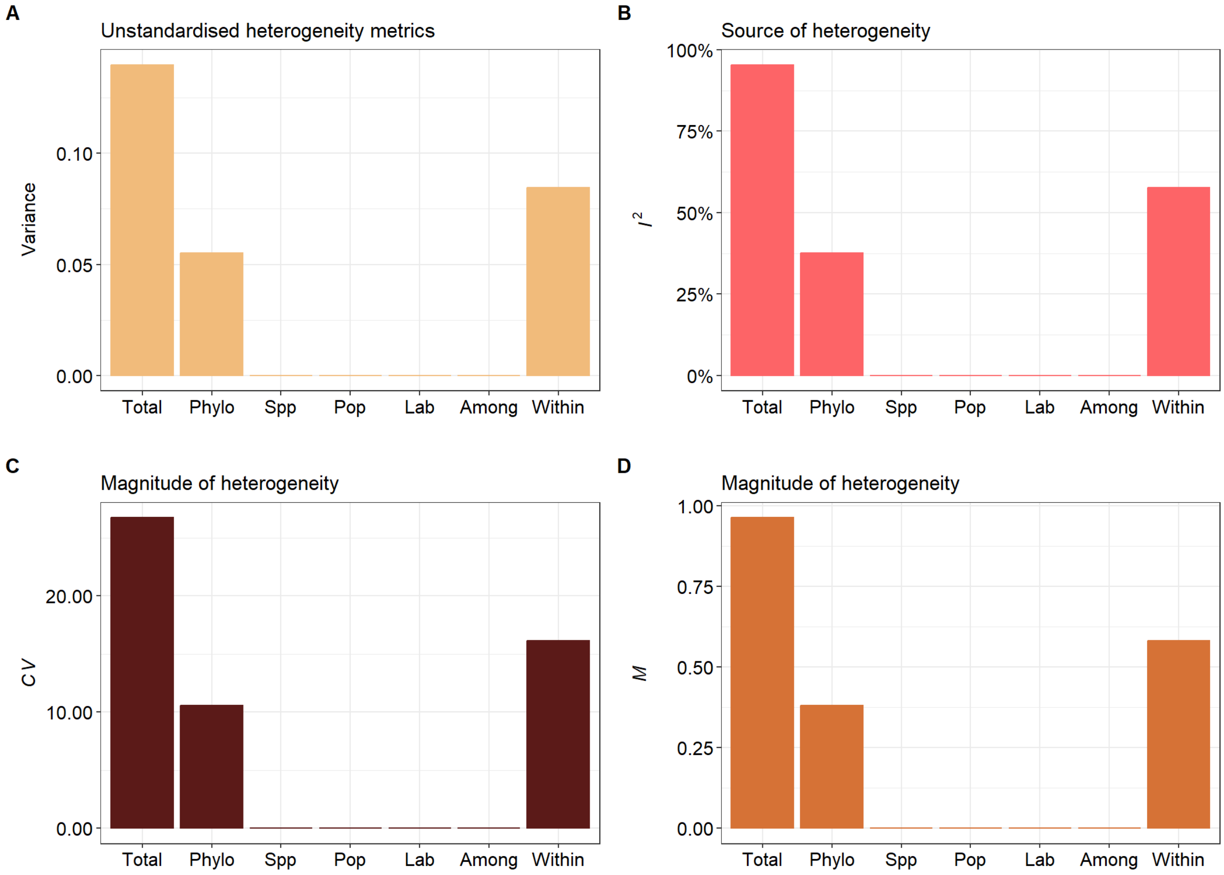

**Supplementary Figure 5.** Effect sizes obtained from the full dataset (considering both maternal and offspring fitness) using a phylogenetic multilevel meta-regression model with 'fitness proxies' as the sole moderator. Orchard plot showing mean estimates (circles with black outlines), 95% confidence intervals (thick whisker), 95% prediction intervals (thin whisker), and individual effect sizes scaled by their precision (coloured circles). k indicates the number of individual effect sizes and, in between brackets, the number of studies. Note that this model was not pre-registered and was conducted after obtaining the results to better understand them.

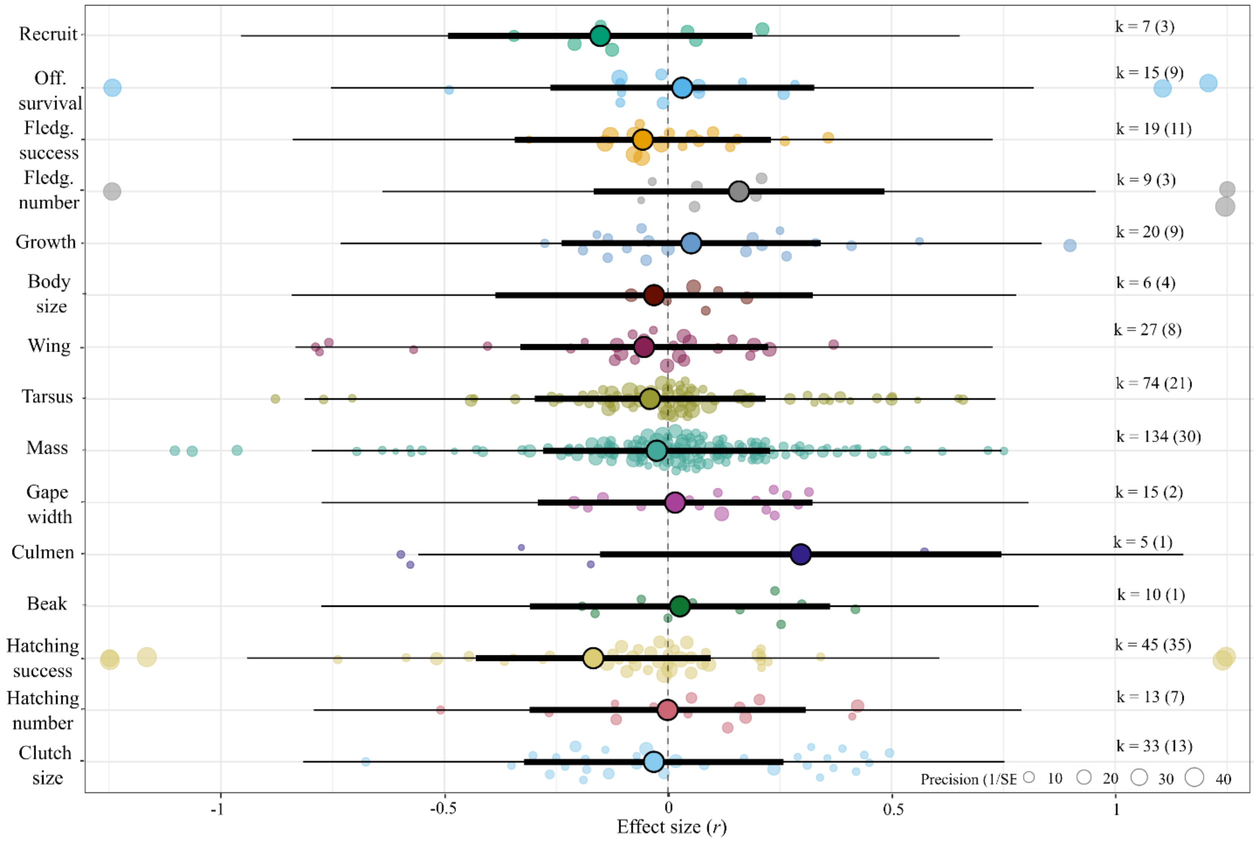
